# Broadly neutralizing antibodies target the coronavirus fusion peptide

**DOI:** 10.1101/2022.04.11.487879

**Authors:** Cherrelle Dacon, Courtney Tucker, Linghang Peng, Chang-Chun D. Lee, Ting-Hui Lin, Meng Yuan, Yu Cong, Lingshu Wang, Lauren Purser, Jazmean K. Williams, Chul-Woo Pyo, Ivan Kosik, Zhe Hu, Ming Zhao, Divya Mohan, Andrew Cooper, Mary Peterson, Jeff Skinner, Saurabh Dixit, Erin Kollins, Louis Huzella, Donna Perry, Russell Byrum, Sanae Lembirik, Yi Zhang, Eun Sung Yang, Man Chen, Kwanyee Leung, Rona S. Weinberg, Amarendra Pegu, Daniel E. Geraghty, Edgar Davidson, Iyadh Douagi, Susan Moir, Jonathan W. Yewdell, Connie Schmaljohn, Peter D. Crompton, Michael R. Holbrook, David Nemazee, John R. Mascola, Ian A. Wilson, Joshua Tan

## Abstract

The potential for future coronavirus outbreaks highlights the need to develop strategies and tools to broadly target this group of pathogens. Here, using an epitope-agnostic approach, we identified six monoclonal antibodies that bound to spike proteins from all seven human-infecting coronaviruses. Epitope mapping revealed that all six antibodies target the conserved fusion peptide region adjacent to the S2’ cleavage site. Two antibodies, COV44-62 and COV44-79, broadly neutralize a range of alpha and beta coronaviruses, including SARS-CoV-2 Omicron subvariants BA.1 and BA.2, albeit with lower potency than RBD-specific antibodies. In crystal structures of Fabs COV44-62 and COV44-79 with the SARS-CoV-2 fusion peptide, the fusion peptide epitope adopts a helical structure and includes the arginine at the S2’ cleavage site. Importantly, COV44-79 limited disease caused by SARS-CoV-2 in a Syrian hamster model. These findings identify the fusion peptide as the target of the broadest neutralizing antibodies in an epitope-agnostic screen, highlighting this site as a candidate for next-generation coronavirus vaccine development.

**One-Sentence Summary:** Rare monoclonal antibodies from COVID-19 convalescent individuals broadly neutralize coronaviruses by targeting the fusion peptide.

## MAIN

Coronaviruses consist of four genera of viruses that infect a range of birds and mammals (*1*). Seven coronaviruses are currently known to cause disease in humans: the alphacoronaviruses HCoV-229E and HCoV-NL63, as well as the betacoronaviruses HCoV-OC43, HCoV-HKU1, SARS-CoV, MERS-CoV and SARS-CoV-2. While the first four coronaviruses are endemic in the human population and generally cause mild disease, the latter three viruses resulted in serious outbreaks in the past 20 years. In particular, the ongoing COVID-19 pandemic by SARS-CoV-2 has caused more than six million deaths since the identification of the virus in late 2019 (*2*). The design of general tools and strategies against SARS-CoV-2 has been complicated by the emergence of variants of concern including the currently dominant Omicron BA.1 and BA.2 variants, which are at least partially resistant to most currently available vaccines and antibody therapeutics (*3–6*). Furthermore, other zoonotic coronaviruses have the potential to spill over from animal hosts and cause infection in humans. For instance, two coronaviruses previously linked only to animal infection, including a more distantly related deltacoronavirus, were recently detected in individuals with flu-like symptoms (*7, 8*). These developments highlight the importance of targeting conserved sites on coronaviruses that are functionally essential and less prone to mutation.

Coronavirus invasion is a multi-step process that involves enzymatic cleavage and major rearrangement of the surface spike protein (*9*). The SARS-CoV-2 spike contains two cleavage sites: a furin cleavage site that is located at the boundary of the S1 and S2 subunits, and an S2’ site that is conserved in all coronaviruses. The spike protein is thought to be cleaved at the S1/S2 site during virus assembly, leaving the S1 and S2 subunits non-covalently linked after virion release from cells. During entry, the SARS-CoV-2 spike protein uses the C-terminal domain (CTD) of the S1 subunit as the receptor-binding domain (RBD) to engage with ACE2 on target cells. Following receptor binding, the S1 subunit is shed and the S2’ site is cleaved by the membrane enzyme TMPRSS2 or endosomal cathepsins (*1*), leading to rearrangement of the S2 subunit to mediate fusion between the virus and cell membranes and insertion of the fusion peptide into the cell membrane.

Much of the protection provided by COVID-19 vaccines is thought to arise from neutralizing antibodies that target the RBD (*10*). Likewise, all currently available therapeutic mAbs target this domain (*3*). While the RBD-ACE2 interaction is essential for SARS-CoV-2 entry, it is only the first of a series of critical steps as described above. The spike elements involved in the latter stages are likely to be more conserved than the RBD, which is capable of retaining or even increasing binding to ACE2 in the face of an array of mutations, even in the ACE2-binding interface (*11*). A stark reminder here is that most COVID-19 therapeutic mAbs currently in the clinic are ineffective against the Omicron variants BA.1 and BA.2 due to extensive RBD mutations (*3, 5*). Therefore, other sites on the spike protein are worth exploring as targets for the development of COVID-19 vaccines and therapeutics that are more resistant to new variants and ultimately for the design of vaccines that target a wider range of coronaviruses. Progress in this direction has been initiated with recent studies identifying several monoclonal antibodies (mAbs) that target the conserved stem helix in the S2 subunit (*12–15*). Nevertheless, to our knowledge, a large-scale survey of the binding landscape of broadly reactive mAbs against coronaviruses has not yet been performed.

### Identification of broadly reactive mAbs from COVID-19 convalescent donors

To identify individuals likely to harbor B cells that produce broadly reactive mAbs, we examined plasma samples of 142 donors from a previously described cohort of SARS-CoV-2 early convalescent individuals in New York (*16*). Using a multiplex bead-based assay, we assessed plasma immunoglobulin G (IgG) reactivity towards spike glycoproteins of the seven human coronaviruses: SARS-CoV-2 (Wuhan Hu-1), SARS-CoV-1, MERS-CoV, HCoV-HKU1, HCoV-OC43, HCoV-NL63 and HCoV-229E. Nineteen donors were selected for mAb isolation and characterization on the basis of their plasma IgG reactivity to SARS-CoV-2 spike and the spike proteins of at least two other betacoronaviruses (Figure. S1A). All 19 selected donors had detectable reactivity to HCoV-OC43 and most had reactivity to HCoV-HKU1 (n= 17), suggesting previous exposure to seasonal betacoronaviruses. However, we did not observe reactivity to the seasonal alphacoronaviruses HCoV-229E and HCoV-NL63 within the detection limits of our plasma screen.

We next interrogated human IgG^+^ memory B cells (MBCs) from the 19 selected donors using a two-stage screening method in which we prioritized isolating mAbs with as great a breadth of reactivity as possible. In the first stage, we screened supernatants from 673,671 stimulated IgG^+^ B cells for binding to the same coronavirus spike panel used for the plasma screen. As expected, SARS-CoV-2 reactive supernatants were frequently observed (20.6% of wells). Supernatants from only 2% (n = 211) of the total MBC culture supernatants screened met our criteria for broad reactivity by binding the SARS-CoV-2 spike and the spike of at least two other non-SARS betacoronaviruses (Figure 1A), highlighting the rarity with which such cross-reactive antibodies arise. For the second stage, we developed a high-throughput, optofluidics assay to isolate individual MBC clones that were actively secreting cross-reactive antibodies using the Berkeley Lights Beacon system (Figure S1B). Candidate cross-reactive MBCs identified in the supernatant screen were pooled and sorted individually into nanoliter-volume pens in an optofluidics chip. Secreted mAbs were then assessed in real-time for binding to beads coated with a cocktail of MERS-CoV and HCoV-OC43 spikes (MERS-CoV/HCoV-OC43^+^), followed by beads coated with SARS-CoV-2 spike (SARS-CoV-2^+^). Double positive MBCs (SARS-CoV-2^+^ and MERS-CoV/HCoV-OC43^+^) were exported off the optofluidics chip for single-cell reverse transcription-PCR (RT-PCR), antibody production as recombinant IgG1, and validation of reactivity to spike protein. In total, we obtained 60 IgG mAbs with reactivity to at least three coronaviruses.

**Figure 1.**
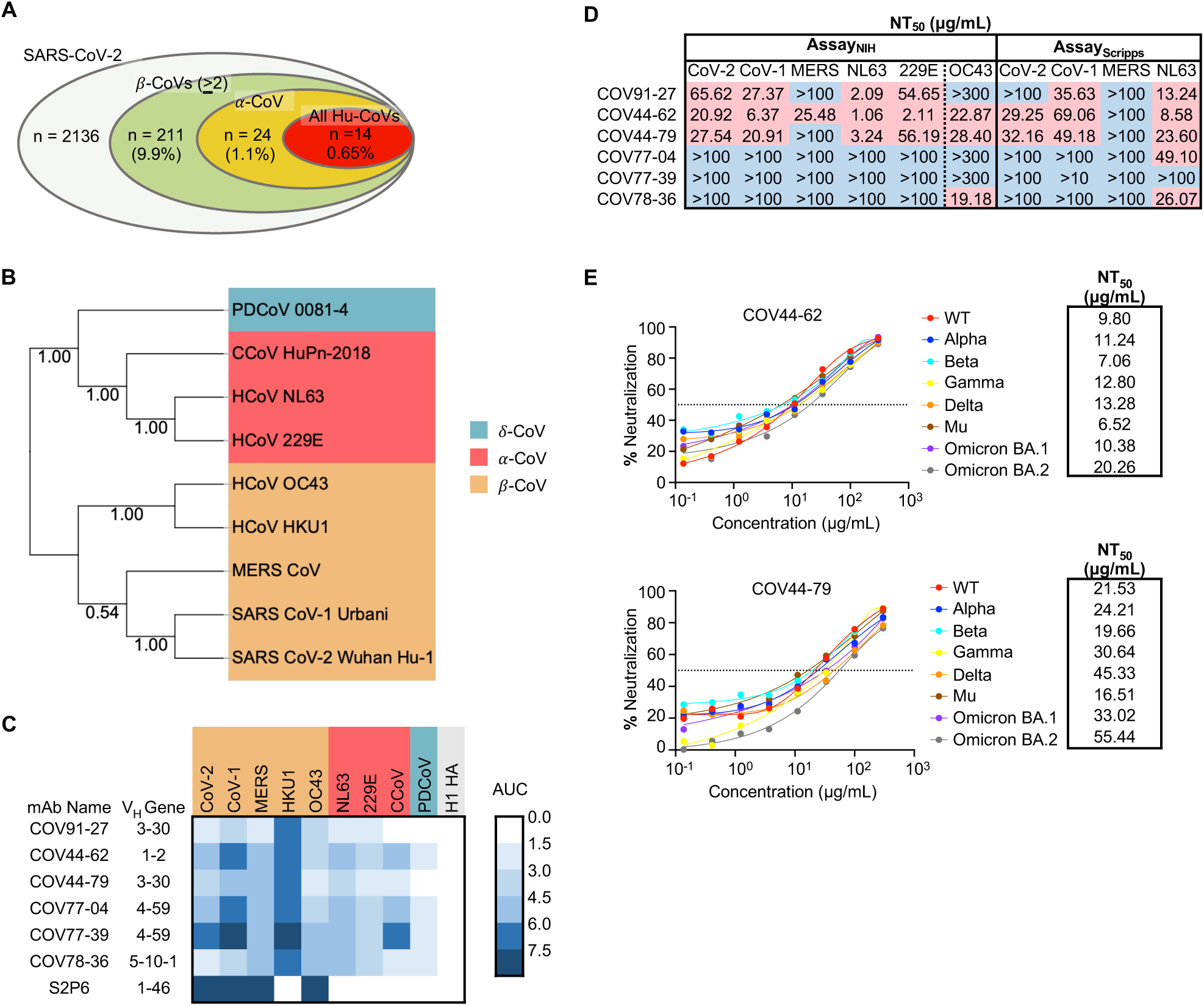
**Broadly neutralizing antibodies target coronaviruses associated with human disease.** (A) Analysis of the frequency of MBCs expressing broadly reactive antibodies from n = 19 donors. Values in parentheses represent the percentage of SARS-CoV-2 reactive supernatants that also bind the specified subsets of non-SARS coronavirus spikes. A total of 10,356 MBC culture supernatants (50-100 B cells/well) was screened. (B) Phylogenetic relationships across the coronavirus spike proteins targeted by the broadly reactive mAbs were inferred by the Neighbor-Joining method in MEGA11 using full-length amino-acid sequences of CoV spike proteins. Bootstrap values from 500 samplings are shown on the branches. (C) Heat map representing the binding of broadly reactive mAbs to spike proteins from coronaviruses across the alpha, beta and deltacoronavirus genera. H1 hemagglutinin was included as a negative control for mAb binding experiments and area under the curve (AUC) values for each antigen are shown after subtraction with values for the negative control antigen CD4. (D) Values represent antibody titer at 50% neutralization (NT_50_) against SARS-CoV-2 Wuhan Hu-1, SARS-CoV-1, MERS-CoV, HCoV-NL63 and HCoV-229E envelope-pseudotyped lentivirus, as well as authentic HCoV-OC43. NT50 values were calculated using the dose-response-inhibition model with 5-parameter Hill slope equation in GraphPad Prism 9. (E) Neutralization of SARS-CoV-2 variants of concern (pseudovirus) by COV44-62 and COV44-79.

To fully interrogate their breadth, we tested the 60 mAbs for binding to spikes from the seven known human coronaviruses. Only six mAbs, COV91-27, COV44-62, COV44-79, COV77-04, COV77-39 and COV78-36, were capable of binding to spike proteins from all seven coronaviruses (Figure 1, B and C). Notably, at least four mAbs in this group also bound to spike from two new coronavirus isolates that have been recently associated with human disease: Canine CoV HuPn-2018 (CCoV-HuPn-2018) and Porcine Deltacoronavirus 0081-4 (PDCoV-0081-4) (*7, 8*) (Figure 1C). The six broadly reactive mAbs were isolated from four different donors, originated from distinct clones, and were encoded by four different VH genes (VH1-2, VH3-30, VH4-59, VH5-10-1). Five of the six mAbs were highly mutated, with VH mutation frequencies ranging from 10 to 13% (Figure S1C). Given that these antibodies were isolated from COVID-19 convalescent individuals in New York approximately one month after the first outbreak in March 2020, the high mutation levels suggest that the B cells were primed during an earlier seasonal coronavirus infection and were possibly reactivated during SARS-CoV-2 infection.

## COV44-62 and COV44-79 broadly neutralize coronaviruses

Next, we assessed the neutralizing potency of the six mAbs against SARS-CoV-2, SARS-CoV-1, MERS-CoV, HCoV-NL63 and HCoV-229E envelope pseudotyped viruses, as well as authentic HCoV-OC43. COV44-62 and COV44-79 showed the broadest functional reactivity, neutralizing the betacoronaviruses SARS-CoV-2, SARS-CoV-1 and HCoV-OC43, as well as the alphacoronavirus HCoV-NL63 and HCoV-229E (Figure 1D and Figure S1D). Moreover, both mAbs were able to neutralize all SARS-CoV-2 variants of concern, including the Omicron BA.1 and BA.2 subvariants, with similar efficacy (Figure 1E). COV44-62 neutralized MERS-CoV in one out of two assays, whereas none of the other mAbs were able to neutralize this virus within the concentrations tested in our assays. COV77-39 was not functional in any of the neutralization assays, despite having the same breadth of binding reactivity as the other five mAbs.

### Broadly reactive mAbs target the coronavirus fusion peptide

To determine the domain of the SARS-CoV-2 spike protein that was targeted by the six broadly reactive mAbs, we assessed mAb binding to the SARS-CoV-2 S2 subunit, as well as the RBD and N-terminal domain (NTD) of the S1 subunit. All six mAbs bound to the S2 subunit, but not the RBD or NTD (Figure 2A). We performed a surface plasmon resonance (SPR) kinetics assay to determine the binding affinity of Fabs derived from these mAbs for pre-fusion stabilized whole SARS-CoV-2 spike (2P, with an intact S1/S2 cleavage site) and the unmodified SARS-CoV-2 S2 subunit. The Fabs bound with low-to-moderate nanomolar affinity to both proteins, but their affinity for the S2 subunit was 3- to 76-fold higher than the corresponding affinity for whole spike (Figure 2B and Figure S2). We did not observe substantial differences between the six Fabs in their binding affinity for the S2 subunit. The mAbs also bound well to a version of S2 that had been stabilized with two proline mutations (S2-2P), but more poorly to a further stabilized version with six proline mutations (S2-HexaPro) (Figure 2C). Next, we performed a competition SPR assay to determine if the six mAbs competed for the same binding site on the S2 subunit. The six mAbs competed with each other for the same site (Figure 2D), but none of them competed with S2P6, a control mAb targeting the stem helix region (*12*), indicating that they bound to a distinct site on the S2 subunit.

**Figure 2.**
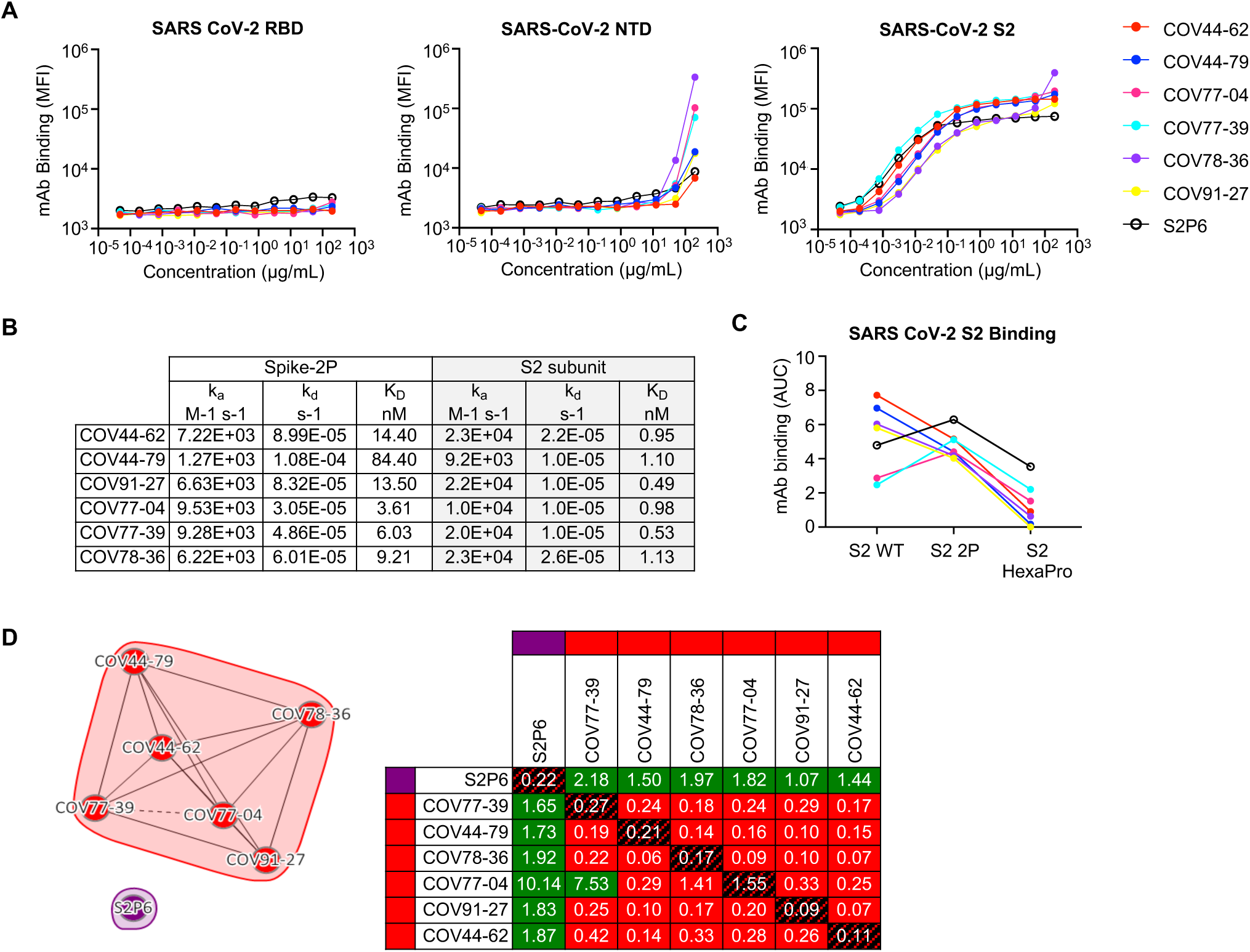
**Broadly reactive mAbs target the same region within the SARS-CoV-2 S2 subunit.** (A) Titration curves for mAb binding to selected regions within the SARS-CoV-2 spike protein: the receptor binding domain (RBD), N-terminal domain (NTD) and the S2 subunit. Interconnected data points are shown without curve fitting. (B) On-rates, off-rates and dissociation constants of the six fusion peptide Fabs for binding to SARS-CoV-2 pre-fusion stabilized spike (2P) with an unmodified furin cleavage site and the non-stabilized S2 subunit. (C) Fusion peptide mAb binding (AUC) to wild-type SARS-CoV-2 S2 subunit and S2 subunit constructs modified with two (2P) or six (HexaPro) stabilizing proline mutations. (D) Epitope binning of broadly reactive antibodies versus the S2 stem-helix targeting mAb S2P6. All included antibodies were tested as both ligands and analytes. Solid lines indicate two-way competition while hashed lines indicate one-way competition. Red boxes indicate competing antibody pairs, green boxes indicate non-competing antibody pairs and hashed filling indicates self-competition.

To further interrogate the specific target site of these mAbs, we subjected them to SPR-based high-throughput peptide mapping using an array of 15-mer linear, overlapping peptides that spanned the length of the entire SARS-CoV-2 S2 subunit (Ser686 to Lys1211, Accession #YP_009724390.1). Mapping analysis revealed that all six mAbs bound to peptides 42-44, which share the _815_RSFIEDLLF_823_ motif (Figure 3A). This motif is located within the SARS-CoV-2 fusion peptide region, directly C-terminal to the S2’ cleavage site. To determine the diversity of this region across coronaviruses, we selected 35 viral isolates representing each of the coronavirus genera – *alpha*, *beta*, *gamma*, and *delta* (Figure 3B). Each amino-acid position in the _815_RSFIEDLLF_823_ motif was conserved in >90% of all viruses selected except F_817_, which was conserved in <50% of all isolates examined (Figure 3, B and C). However, variability at this position was still limited where ∼ 86% of the isolates had a phenylalanine or alanine, indicating a degree of conservation (Figure 3B). The fusion peptide appears partially surface-exposed in a range of coronavirus spike proteins, including SARS-CoV-2, SARS-CoV, MERS-CoV and MHV (*17, 18*) (Figure 3C and Figure S3A). However, antibody access to this site may be partially occluded by the S1 subunit on an adjacent protomer, which is consistent with the stronger binding displayed by the mAbs for the S2 subunit relative to the pre-fusion-stabilized SARS-CoV-2 spike (Figure 2B and Figure S3B).

**Figure 3.**
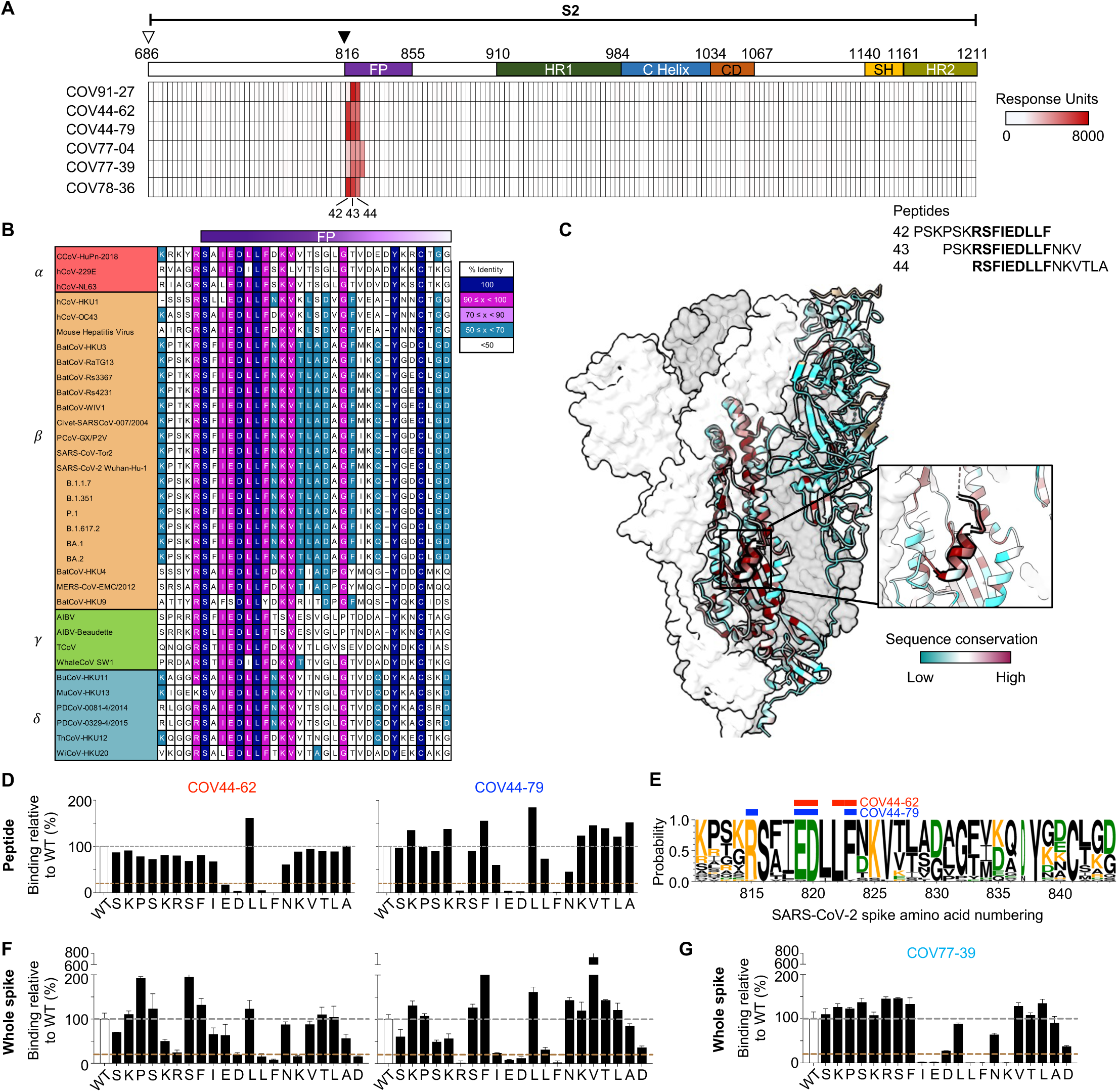
**Broadly neutralizing antibodies target the conserved fusion peptide.** (A) Heat map of SARS-CoV-2 S2 peptide array. Binding responses were assessed by SPR using a 15-mer peptide array with 12 aa overlay covering the entire S2 subunit. Each column within the map represents a single peptide. The white triangle shows the S1/S2 cleavage site and the black triangle shows the S2’ cleavage site. FP, fusion peptide; HR1, heptad repeat 1; C helix, central helix; CD, connector domain; SH, stem helix; HR2, heptad repeat 2. (B) Sequence alignment of the fusion peptide from 34 viral isolates representing a diverse group of coronaviruses across four genera. Performed using MAFFT v7.450 using a BLOSUM62 scoring matrix and the L-INS-I algorithm. (C) Sequence conservation of pre-fusion SARS-CoV-2 spike protein (PDB 6VSB) with the fusion peptide (aa 816-843) highlighted in black. Inset shows a magnified view of this region. (D) Alanine scan evaluating the binding of COV44-62 and COV44-79 to the SARS-CoV2 fusion peptide. Responses were normalized to the wild-type sequence. A cut-off of 20% (brown hashed line) was used to identify residues that were critical for binding. (E) Sequence logo plot of diversity within the fusion peptide region of coronaviruses from 35 isolates, built using WebLogo 3. Height is proportional to the probability of an amino acid at a given position and amino-acid residues are colored by charge. Narrow stacks (amino acids) indicate deletions or gaps in the sequences. Numbering is based on the SARS-CoV-2 Wuhan-Hu-1 sequence. The key residues in the epitope footprints of mAbs COV44-62 (red) and COV44-79 (blue), based on peptide alanine scanning, are highlighted above the logo plot. (F-G) Amino acids critical for the binding of COV44-62, COV44-79 and COV77-39 identified by shotgun alanine mutagenesis of all residues on the S2 subunit. Key residues were identified based on a <20% signal relative to wild-type spike (brown hashed line), with no corresponding loss of signal for a control mAb, which targets the spike protein but does not bind to this site (see Figure S3C). The error bars show half the range (max – min signal).

The finding that these mAbs target the fusion peptide is consistent with their reduced binding to the HexaPro S2 construct (Figure 2C), which includes a non-conservative F817P mutation in the middle of this site (*19*). To identify the specific amino acids that are important for binding of the fusion peptide-specific mAbs, we performed a peptide alanine scan on this region and focused on analyzing the residues targeted by the broadly neutralizing mAbs COV44-62 and COV44-79 (Figure 3D). Four amino acids, E819, D820, L822 and F823, were crucial (<20% binding to Ala mutant) for binding of COV44-62, where mutation of the F823 residue completely abolished binding. Similarly, the binding site of COV44-79 encompassed E819, D820 and F823, but also included R815, the site of S2’ cleavage, as a critical residue (Figure 3D). All five epitope residues that were identified as important for COV44-62 or COV44-79 binding are among the most conserved residues in the coronavirus spike protein. D820 and L822 are completely conserved, while only one out of the 35 coronaviruses examined has a different amino acid at the 815, 819 and 823 positions (Figure 3, B and E). Amino-acid mutations at the peptide level may have slightly different effects from mutations in the intact spike protein, where modified interactions with surrounding S2 residues may also affect antibody binding. Therefore, we also analyzed the residues that were critical for mAb binding in the context of the whole spike protein using a shotgun alanine mutagenesis approach. In this assay, every amino acid in the S2 subunit of intact spike was individually mutated to generate a panel of spike mutants for screening with COV44-62 and COV44-79, as well as a non-neutralizing fusion peptide-specific mAb, COV77-39 (Figure 3, F and G, and Figure S3, C and D). In general, this assay identified a greater number of residues as important for mAb binding, including some with a more intermediate phenotype. For COV44-62, D820, L822 and F823 were once again crucial for binding, while K825 and D830, as well as R815, were also identified as important (Figure 3F). For COV44-79, the results closely matched the peptide alanine scan, with the same four amino acids, R815, E819, D820 and F823, identified as the most critical. Interestingly, the non-neutralizing mAb COV77-39, which has similar binding affinity for the SARS-CoV-2 spike and S2 subunit (Figure 2B), targeted a group of residues that did not include R815 (Figure 3G), suggesting that binding to the S2’ cleavage site may be a distinguishing property of neutralizing mAbs against this site.

### Crystal structures of anti-fusion peptide antibodies

To elucidate the molecular characteristics of anti-fusion peptide antibodies that neutralize SARS-CoV-2, the Fabs of COV44-62, COV44-79, and COV91-27 were complexed with 15-mer peptides containing the fusion peptide sequence (Figure 4). Crystal structures were determined to 1.46 Å, 2.8 Å and 2.3 Å resolution, respectively (Figure 4, Figure S4 and table S1). Fourteen of the 15 peptide residues were visible in the electron density map for COV44-62 (Figure S4A), 13 of which have a buried surface area (BSA) > 0 Å^2^ in complex with antibody. For COV44-79, 12 of the 15 peptide residues were visible in the electron density map (Figure S4A), and 10 of which have a buried surface > 0 Å^2^. Similarly, in COV91-27, 12 peptide residues were visible (Figure 4C and Figure S4A), with nine exhibiting a buried surface > 0 Å^2^. The fusion peptide forms a helix as in the prefusion state of the SARS-CoV-2 spike (Figure S4B). All three complementarity-determining regions (CDRs) of the heavy chain (HC) of all three Fabs are involved in peptide recognition whereas CDR1 and CDR3 of the light chain (LC) of COV44-62 and only LCDR3 of COV44-79 and COV91-27 contact the peptide (Figure 4, A to C). The BSA on the each Fab is dominated by the heavy chain and is 791 Å^2^ for COV44-62 (627 Å^2^ by HC and 164 Å^2^ by LC), 573 Å^2^ for COV44-79 (505 Å^2^ by HC and 129 Å^2^ by LC), and 573 Å^2^ for COV91-27 (447 Å^2^ by HC and 126 Å^2^ by LC). The fusion peptide makes side-chain and backbone H-bonds and salt bridges with COV44-62 mainly through K814, R815, E819, D820, L822, F823 and N824, and hydrophobic interactions via I818, L822, and F823 (Figure 4D). These residues include the key residues, E819, D820, L822 and F823 identified by site-directed mutagenesis (Figure 3D). The fusion peptide did not form as many interactions with COV44-79 and COV91-27 (Figure 4, E and F). However, R815, E819, and D820 contributed H-bonds and salt bridges, and I818, L822, and F823 made hydrophobic interactions. In all three antibodies, R815, S816, I818, E819, D820, L822, and F823 contributed the most buried surface area to the interaction (Figure 4, G to I). The structural results are consistent with the mutagenesis data with peptide and spike protein that identify the key binding residues (Figure 3, D to F, and Figure S3D). Importantly, the arginine at the S2’ cleavage site is also involved in recognition by these anti-fusion peptide antibodies. However, the approach angle of the three Fabs to the fusion peptide differs from each other, although the antibodies all interact with one face of the fusion peptide helical structure (Figure S4D). Superimposition of the fusion peptide structures onto an intact SARS-CoV-2 spike trimer structure in the pre-fusion state showed a potential clash with the S protein, suggesting a conformational change or conformational dynamics around the fusion peptide is required to accommodate antibody targeting of the fusion peptide (Figure S4C). Notwithstanding, these antibodies have neutralization activity against SARS-CoV-2 and therefore must be able to interact with the fusion peptide on the virus (Figure 1D).

**Figure 4.**
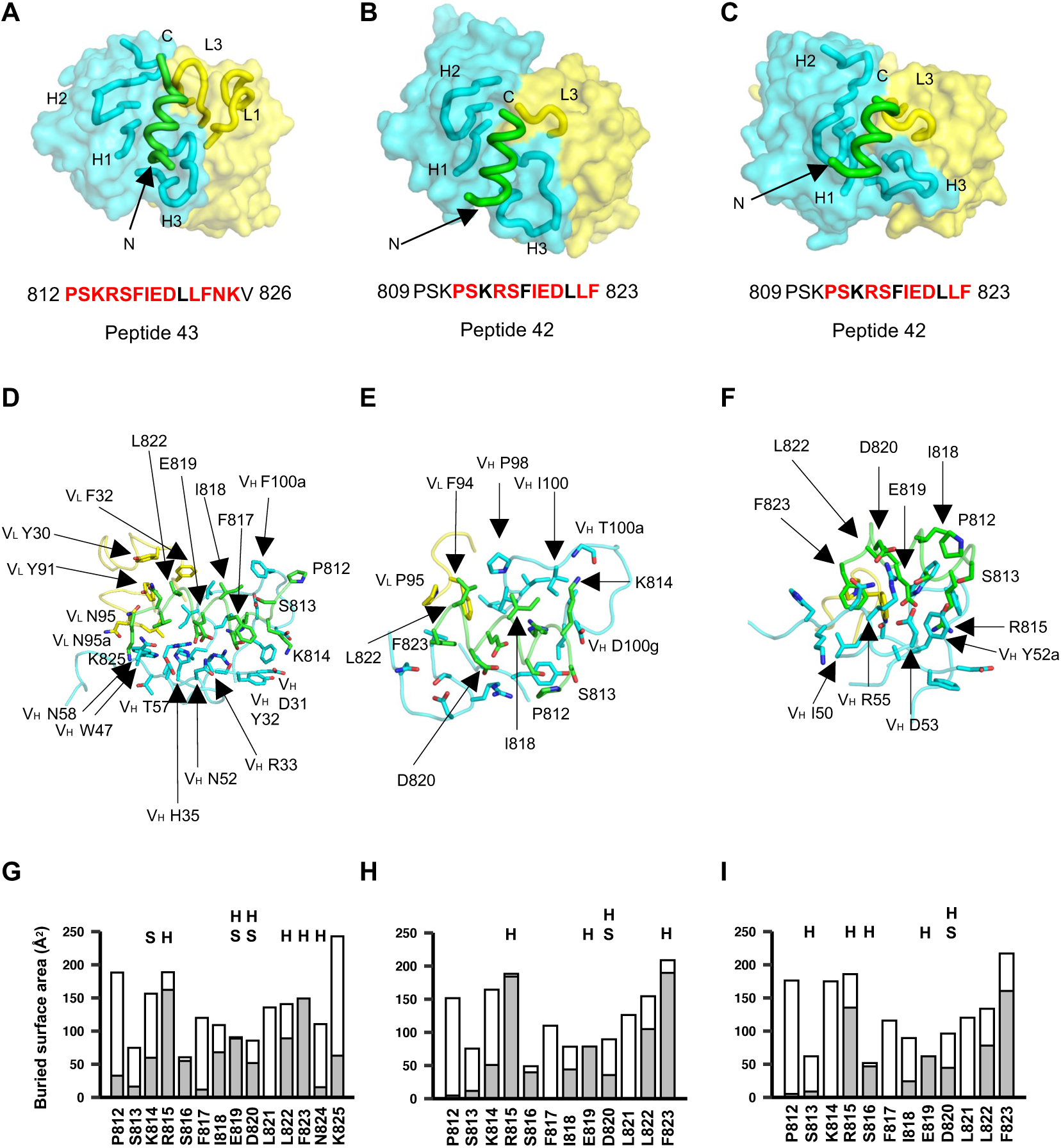
**Crystal structures of COV44-62, COV44-79, and COV91-27 in complex with SARS-CoV-2 fusion peptide.** (A-C) The overall interactions of (A) COV44-62, (B) COV44-79, and (C) COV91-27 with the fusion peptide. Fabs are shown in a molecular surface and the CDRs and peptides are represented as tubes. Cyan and yellow represent the heavy and light chains of the Fabs. Peptides are shown in green. H1, H2, H3, L1, and L3 denotes CDRs in the heavy (H) and light (L) chains. The resolution of the crystal structures are 1.46 Å, 2.8 Å, and 2.3 Å for the COV44-62, COV44-79, and COV91-27 complexes. Peptide residues observed in the crystal structure are in bold and residues involved in interaction with antibody (buried surface area >0 Å^2^) are in red. (D-F) Details of the interactions between (D) COV44-62, (E) COV44-79, and (F) COV91-27 with the fusion peptide. VH and VL indicate the variable domains of the heavy (H) and light (L) chains. Kabat numbering was used for the Fabs and numbering in the native spike protein for the fusion peptide. The colors for the heavy chain, light chain, and fusion peptide are as in (A). (G-I) Buried surface area (in gray) and accessible surface area (in white) of each residue of the fusion peptide in complex with antibody are shown in the stacked bar chart. Residues that form polar interactions with COV44-62, COV44-79, and COV91-27 are denoted with “H” if they form a hydrogen-bond or “S” for a salt bridge on top of the corresponding bar. Buried and accessible surface areas were calculated with PISA ((*31*)).

### Response to the fusion peptide following COVID-19 vaccination and infection

To investigate the antibody response to the fusion peptide after infection and vaccination, we compared the binding of polyclonal IgG from mRNA-1273-vaccinated donors (Figure S5A), COVID-19 convalescent individuals, and COVID-19-naïve individuals whose samples were collected prior to the pandemic. mRNA-1273 encodes spike protein that has an intact S1/S2 cleavage site but is pre-fusion stabilized with two proline mutations, and thus may elicit a different antibody response to the fusion peptide compared to that generated during natural infection. Polyclonal IgG from these donors was purified from human sera or plasma, normalized to equal concentrations and tested for binding to the SARS-CoV-2 fusion peptide (peptide 43) (Figure S5B). All 13 COVID-19-naïve donors had minimal binding to both peptides, indicating a minor contribution by potential previous seasonal coronavirus infections to circulating fusion peptide-specific IgG. While there was an increased response in several vaccinees after the 2^nd^ vaccine dose, most individuals did not produce specific IgG above the baseline one month after this dose (*P* > 0.999 vs baseline), and this was not enhanced by the administration of a booster. As a group, the COVID-19 convalescent donors did not have significantly higher responses to the fusion peptide compared to the vaccinated donors (*P* > 0.999 versus post-2^nd^ dose). However, several convalescent donors in this group had the highest responses in all three cohorts, suggesting that natural SARS-CoV-2 infection can trigger a strong antibody response to the fusion peptide in specific individuals.

### COV44-62 and COV44-79 inhibit membrane fusion

Given the binding of COV44-62 and COV44-79 to the fusion peptide, we hypothesized that they would inhibit spike-mediated cell fusion, which relies on insertion of the fusion peptide into the target cell membrane. We observed that COV44-62 and COV44-79 inhibited the fusion of cells expressing SARS-CoV-2 spike and cells expressing the ACE2 receptor (Figure 5A).

**Figure 5.**
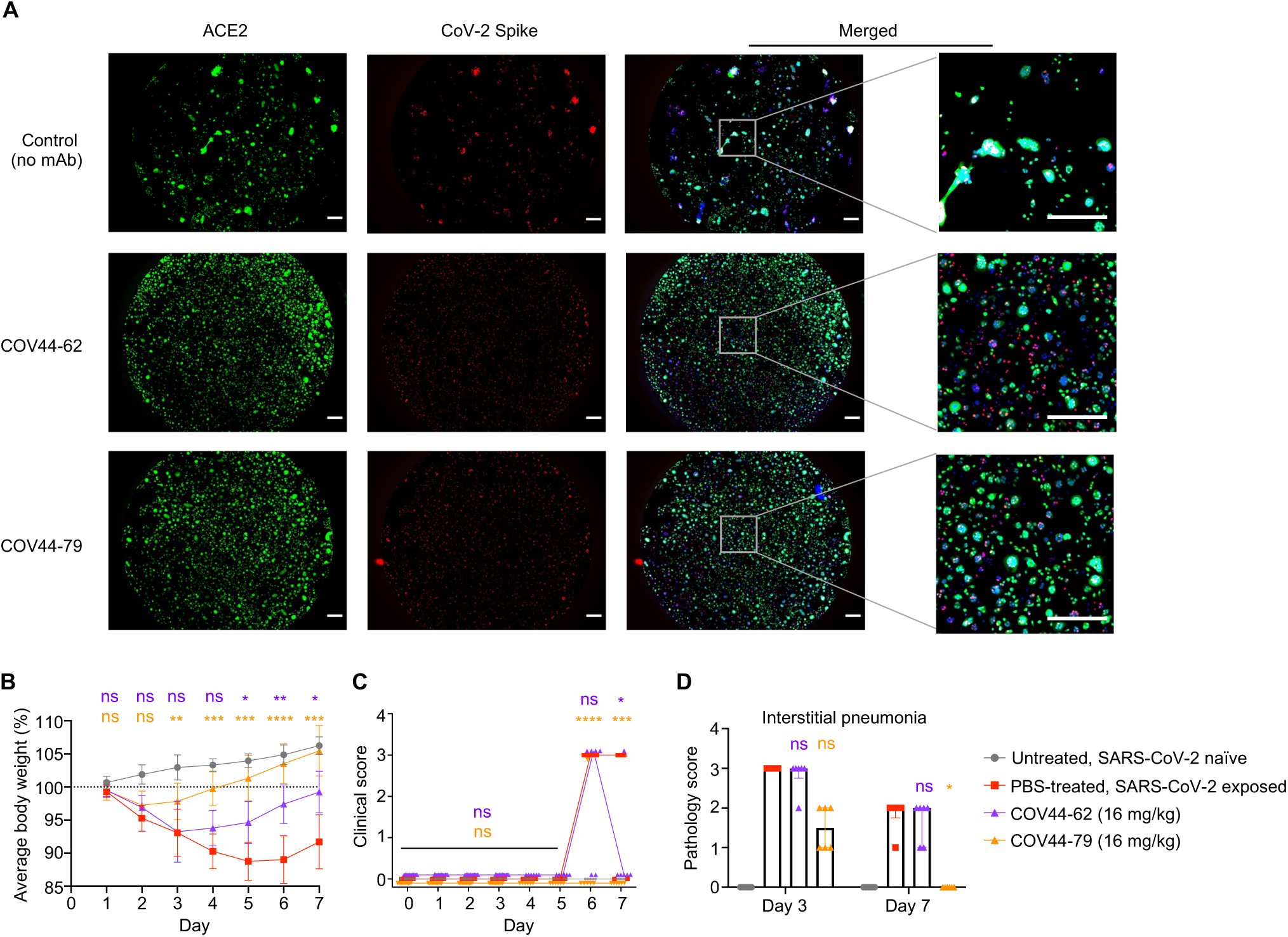
**COV44-62 and COV44-79 inhibit SARS-CoV-2 spike-mediated fusion and COV44-79 limits disease in a Syrian hamster model.** (A) Images of fusion between HeLa cells stably expressing SARS-CoV-2 spike (RFP) and HeLa cells stably expressing the ACE2 receptor (GFP) after counter-staining with Hoechst (blue). Cells were co-cultured in the presence of COV44-62, COV44-79 or without a mAb (control). Scale bar, 500 μm. (B) Weight change for SARS-CoV-2 naïve animals versus virus-exposed animals that were mock-treated or treated with 16 mg/kg of mAb. Statistical significance for average body weight was analyzed across the 7-day time-course using a mixed-effects repeated measures model with Dunnett’s post-test multiple comparison (n=12 animals from Day 0-6 and n=6 animals from Day 7-12). Error bars show mean ± SD. (C and D) Clinical scores (C) and pathology scores (D) for SARS-CoV-2 naïve animals versus virus-exposed animals that were mock-treated or treated with 16 mg/kg of mAb. Statistical analyses for clinical (Days 0-7) and pathology (Days 3 and 7) scores were analyzed by a Kruskal-Wallis test with Dunn’s post-test multiple comparison (n=6-12 animals per condition), between the mAb-treated and mock-treated groups on each day. In (C), lines connect the median values. In (D), error bars show the median ± interquartile range.

### COV44-79 limits disease in the Syrian hamster model

We evaluated the *in vivo* efficacy of COV44-62 and COV44-79 against SARS-CoV-2 infection in the Syrian hamster model, a well-established model recapitulating features of moderate to severe COVID-19 in humans (*20–22*). For this assay, we converted the Fc regions of the two mAbs to hamster IgG2 to allow optimal Fc function. The mAbs were administrated intraperitonially at 16 mg/kg, followed by intranasal administration of 5 log10 PFU of SARS-CoV-2 WA01 24 hours later (Figure 5, B to D, and Figure S6). Change in body weight and a blinded clinical score were noted daily for clinical assessment. Consistent with previous findings, hamsters from PBS-treated controls had a more than 10% body weight reduction through day 6 post-virus exposure, followed by a rebound in weight, whereas mock-exposed hamsters continue to gain weight throughout the study period (*16*) (Figure 5B). Hamsters treated with COV44-79, and to a lesser extent COV44- 62, had a smaller decrease in body weight and recovered more quickly than untreated hamsters (*P* < 0.01 from days 3-7 for COV44-79 and *P* < 0.05 from days 5-7 for COV44-62) (Figure 5B). A similar trend was found when the clinical scores were analyzed (Figure 5C). Rapid breathing was observed in all six hamsters in the sham-treated SARS-CoV-2–exposed group starting on day 6, with two hamsters recovering on the following day. Four of six hamsters from the COV44-62- treated group developed clinical signs by day 6 and one hamster retained signs on Day 7 (*P* < 0.05 on day 7) (Figure 5C). In contrast, only one hamster treated with COV44-79 had a rapid respiratory rate on day 6 and recovered the next day (*P* < 0.001 on days 6 and 7). Furthermore, semiquantitative scoring of interstitial pneumonia revealed that hamsters treated with COV44-79 had lower pneumonia than untreated hamsters on day 7 (P < 0.05) (Figure 5D).

## Discussion

In this study, we identified six mAbs that bound to spike protein from all human alpha- and beta- coronaviruses, with two mAbs neutralizing at least six out of seven viruses. These mAbs target the fusion peptide region, which plays a critical role during coronavirus invasion, has an identical sequence in all SARS-CoV-2 variants of concern, and is highly conserved across the four coronavirus genera. Moreover, this region is at least partially exposed on the surface of the pre- fusion spike of various coronaviruses including SARS-CoV-2, SARS-CoV, MERS-CoV and MHV (*17, 18*). The broad conservation and functional importance of the fusion peptide highlight the potential of this site as a candidate for coronavirus vaccine development. These findings have parallels to work done on HIV-1 gp120, where the surface-exposed fusion peptide was identified as a target of neutralizing mAbs (*23*). This discovery led to the investigation of the HIV-1 fusion peptide as an immunogen to elicit broadly neutralizing antibodies in immunized animals (*24*), and subsequent animal studies have increased the potency and breadth of the fusion peptide-targeting antibody response (*25*).

Despite these potential advantages, the coronavirus fusion peptide has not been a major focus of attention for development of therapeutic mAbs and COVID-19 vaccines. The main drawback of the mAbs described here is their comparatively low *in vitro* neutralization potency that may be due to their relatively weak binding to intact spike, which is enhanced only when the S1 cap is removed. These mAbs fit into a wider trend of a trade-off between potency and breadth: highly potent mAbs targeting the RBD are restricted to sarbecoviruses and most do not neutralize all variants of SARS-CoV-2 (*3, 5, 6, 16, 26*), while mAbs targeting the stem helix (*12–15*) and those identified here have greater breadth but are less potent. However, as previously reported for some of the anti-stem helix mAbs, COV44-79 performed better in the hamster model than expected, suggesting that it may function in a way that is not captured efficiently in the neutralization assay, including involving effector functions through the Fc region. Moreover, there is substantial scope for improvement for mAbs with this specificity and subsequent studies may discover mAbs with higher potency, as was seen with the HIV-1 fusion peptide (*27*). *In vitro* and *in vivo* techniques have recently been described to improve antibody affinity and potency and may be useful here (*28, 29*).

Additionally, vaccination with fusion peptide constructs may trigger a polyclonal response of greater magnitude and potency against this site. Here, we found that three doses of the mRNA- 1273 vaccine did not produce strong antibody responses to the fusion peptide. While the response in COVID-19 convalescent individuals was variable, several individuals had strong antibody responses after SARS-CoV-2 infection, potentially due to the activation of fusion peptide-specific memory B cells primed during a previous seasonal coronavirus infection. This observation is consistent with greater exposure of the S2 subunit to B cells during natural infection due to the uncoupling of the S1 subunit, an event that likely occurs less frequently with pre-fusion stabilized spike protein. Of note, a recent study reported that depleting fusion peptide-specific antibodies from the serum of COVID-19 convalescent patients resulted in a 20% reduction in SARS-CoV-2 neutralization, indicating that polyclonal antibodies targeting this site play an appreciable role (*30*). A nanoparticle vaccine based on the fusion peptide may be even more effective than natural infection in triggering specific antibody responses and should be relatively straightforward to design given the linearity of the target epitope.

The COVID-19 pandemic has shown that it is difficult to predict the emergence of novel coronaviruses or variants and act accordingly to prepare specific tools in advance. Therefore, the development of wider strategies would be invaluable to enhance pandemic-preparedness efforts for targeting of coronaviruses and other viruses, such as influenza virus. This study offers proof-of-principle that mAbs that target the highly conserved fusion peptide region can neutralize coronaviruses from different genera and limit coronavirus-caused disease *in vivo*, warranting further investigation of this site as a candidate for a broader coronavirus vaccine.

## Acknowledgments

We thank the blood sample donors at the New York Blood Center; Sandhya Bangaru, Gabriel Ozorowski, Alba Torrents de la Peña, and Andrew Ward for providing HCoV-OC43, MERS-CoV and H1 HA; Gavin Wright and Nicole Muller-Sienerth (Wright lab, University of York) for providing recombinant CD4; Melanie Cohen and Julie Laux for assistance with cell sorting. We thank Nick Vaughan, Kurt Cooper, Becky Reeder, Marisa St Claire, Kyra Hadley, David Drawbaugh, Amanda Hischak, Randy Hart, and Nejra Isic for assistance with hamster experiments. This work is supported by the Division of Intramural Research and the Vaccine Research Center, National Institute of Allergy and Infectious Diseases (NIAID), National Institutes of Health (NIH). (ID, SM, JWY, PDC, MRH, JRM, JT). This project has been funded in whole or in part with Federal funds from the National Institute of Allergy and Infectious Diseases (NIAID), National Institutes of Health (NIH), U.S. Department of Health and Human Services (DHHS) under Contract No. HHSN272201800013C. NIH grant R01AI132317 (DN and LP), HHSN contract 75N93019C00073 (JKW and ED), and the Bill and Melinda Gates Foundation grant INV-004923 (IAW)

## Author contributions

Conceptualization: JT, CD. Methodology: CD, CT, LPe, C-CDL, THL, MY, YC, LW, IK, ZH, RW, AP, ED, DG, ID, SM. Formal analysis: CD, CT, LPe, MY, YC, LW, C-CDL, THL, C-WP, JS, ED. Data curation: MP. Investigation: CD, CT, LPe, MY, LW, C-CDL, LPu, C-WP, JKW, THL, MZ, MP, DM, AC, SD, EK, LH, DP, SL, YZ, ESY, MC, KL. Resources: RW, SM. Writing - original draft: CD, CT, MY, JT. Writing - review & editing: all authors. Visualization: CD, CT, LPe, MY, YC, LW, C-CDL, JKW, ED, JT. Supervision: ED, DG, SM, JWY, CS, PDC, DN, MRH, JRM, IAW, JT. Funding acquisition: CS, PDC, DN, MRH, JRM, IAW, JT.

## Data and materials availability

All data associated with this manuscript are available in the main text or the supplementary materials. Crystal structures will be deposited into the Protein Data Bank. Materials described in this manuscript will be available through a material transfer agreement (MTA) with the National Institute of Allergy and Infectious Diseases.

---

**Figure S1.**
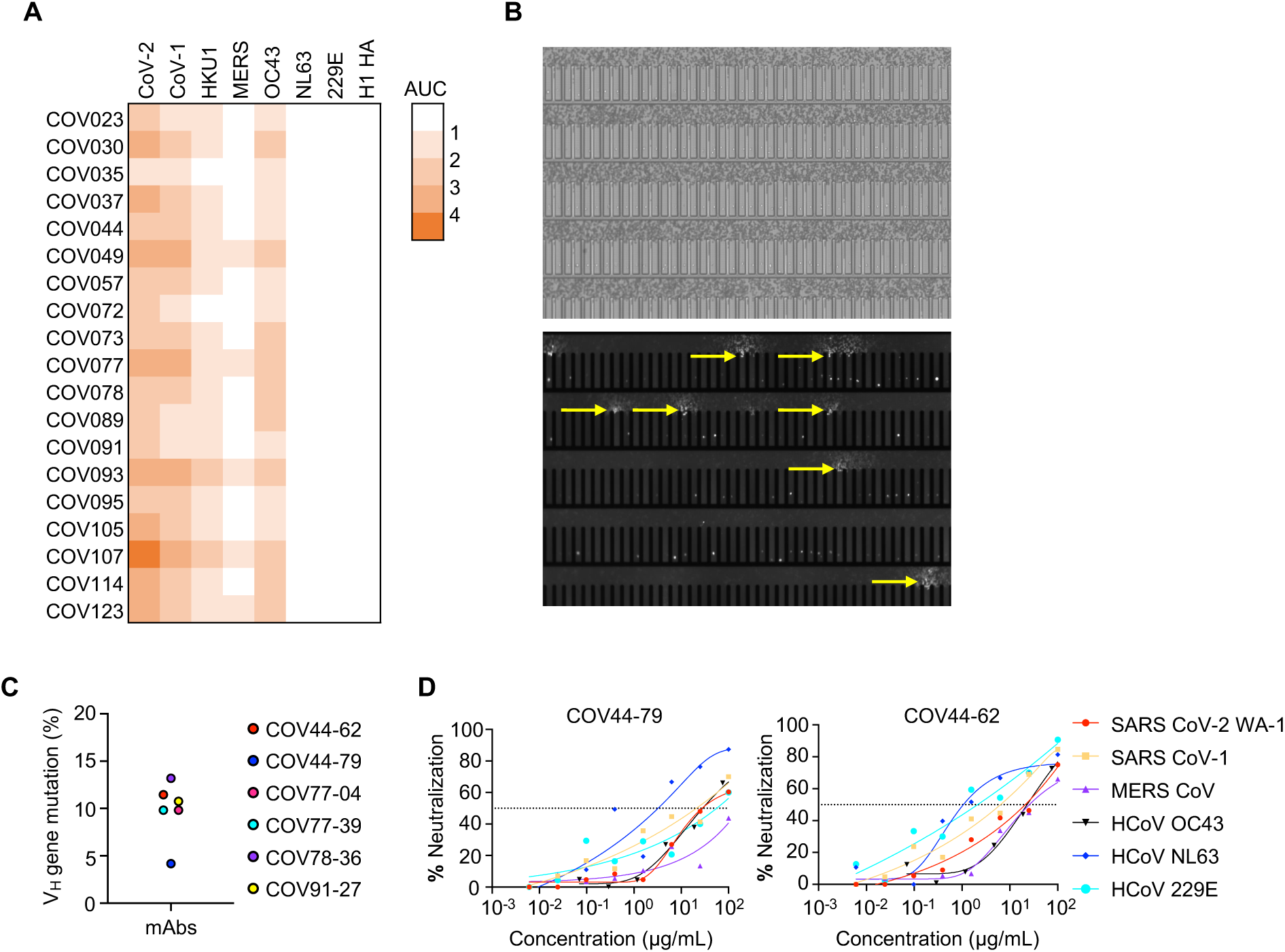
**COVID-19 convalescent donor screening and isolation of broadly neutralizing mAbs. (A**) COVID-19 convalescent plasma reactivity to the seven human coronaviruses. The heat map represents area under the curve (AUC) values for each antigen after subtraction with values for the negative control antigen CD4. H1 haemagglutinin was included as a control. (**B**) Representative images show optofluidic screening of MBC secreted antibodies for cross-reactivity. The top panel shows individual MBCs sorted into nanopens and the bottom panel shows the fluorescent signal of antigen-specific antibodies binding to coronavirus spike-coated beads (yellow arrows). (**C**) VH gene mutation levels of fusion peptide-targeting mAbs. (**D**) Neutralization curves of COV44-62 and COV44-79 against authentic HCoV-OC43, as well as SARS-CoV-2 Wuhan Hu-1, SARS-CoV-1, MERS-CoV, HCoV-NL63 and HCoV-229E envelope-pseudotyped virus.

**Figure S2:**
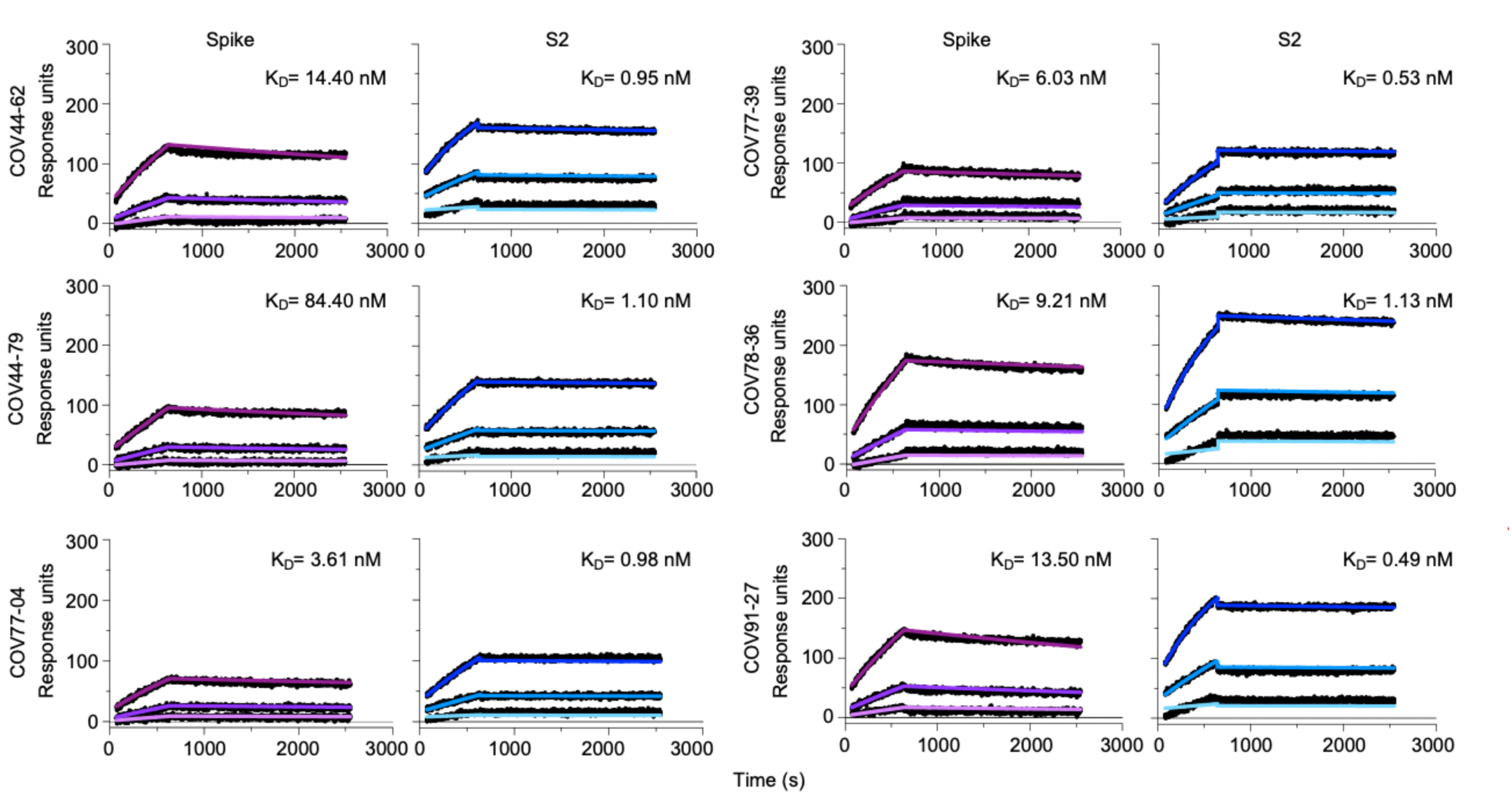
Kinetics curves for binding of fusion peptide Fabs to SARS-CoV-2 spike and S2 subunit. Curves are shown for binding to SARS-CoV-2 pre-fusion stabilized spike (2P) with an unmodified furin cleavage site and the non-pre-fusion stabilized S2 subunit. Black dots show raw data points and blue and purple lines show fitted curves.

**Figure S3.**
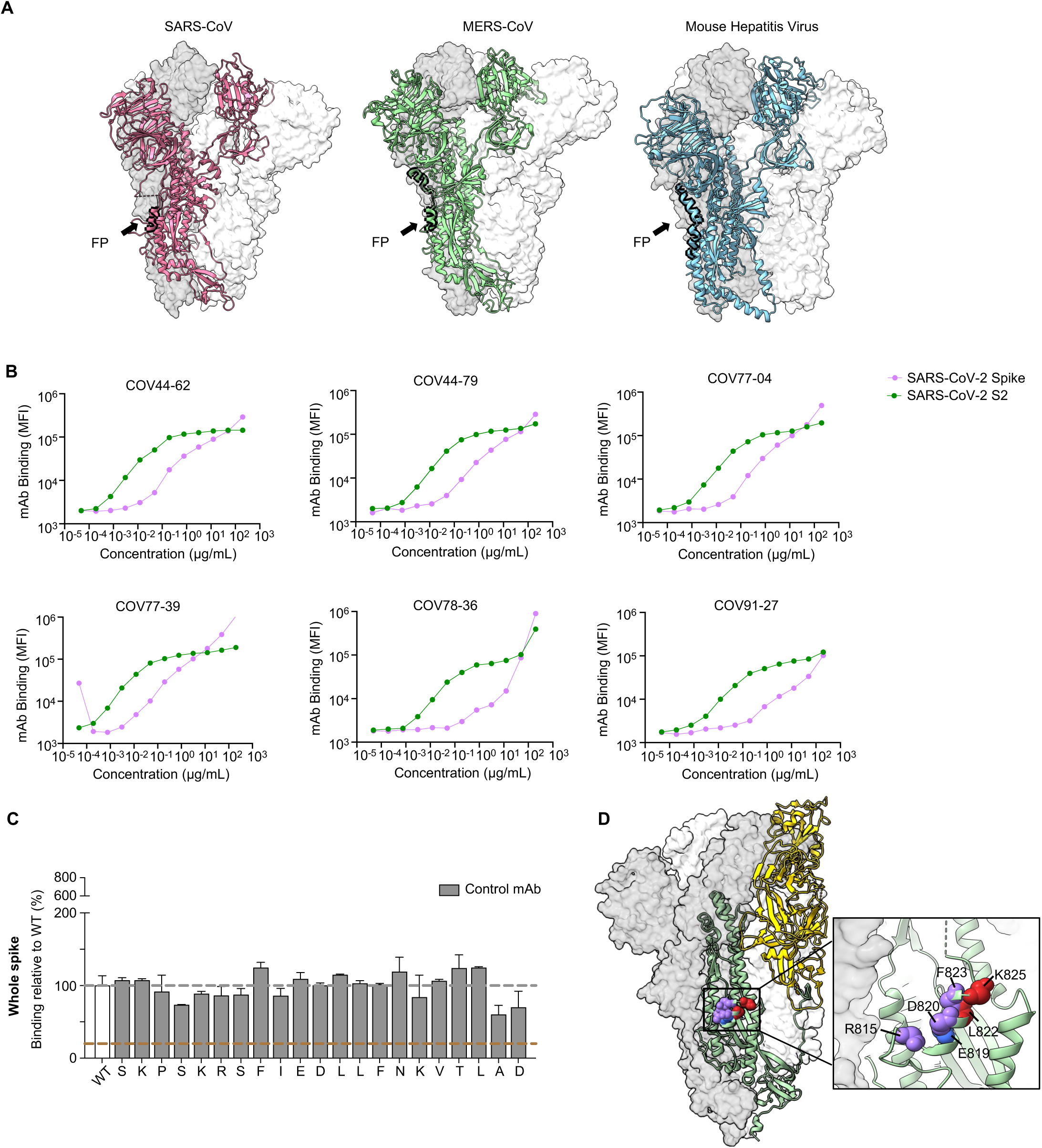
Binding profile of broadly reactive mAbs targeting the fusion peptides. (**A**) Surface exposure of fusion peptide loops (highlighted in black) of SARS-CoV-1 (PDB ID 5X58), MERS-CoV- England 1 (PDB ID 6Q04), Mouse Hepatitis Virus (PDB ID 3JCL) amino-acid residues 870-897. The fusion peptide loop for each virus is highlighted in black corresponding to residues 870-897, 888-915, and 798-825, respectively. (**B**) Titration curves of broadly reactive mAb binding to SARS-CoV-2 spike trimer (2P, furin cleavage site intact) and unmodified SARS-CoV-2 S2 subunit monomer in a bead-based assay. Interconnected data points are shown without curve fitting. (**C**) Amino acids important for the binding of a negative control mAb identified by shotgun alanine mutagenesis of all residues on the S2 subunit. The control mAb binds to the spike protein but not to this region of the protein. Any residues where the binding of the control was <20% relative to wild-type spike was not considered a target residue of COV44-62, COV44-79 or COV77-39. Any alanines in the sequence were mutated to serines. The error bars show half the range (max – min signal). (**D**) Residues important for the binding of COV44-62 and COV44-79, based on shotgun mutagenesis, are shown on the SARS-CoV-2 spike structure. Residues important for COV44- 62 binding are in red, those important for COV44-79 are in blue, and those important for both are in purple. D830, which was identified as a target residue of COV44-62, was not resolved in this structure (PDB 6VSB) and is not shown here.

**Figure S4.**
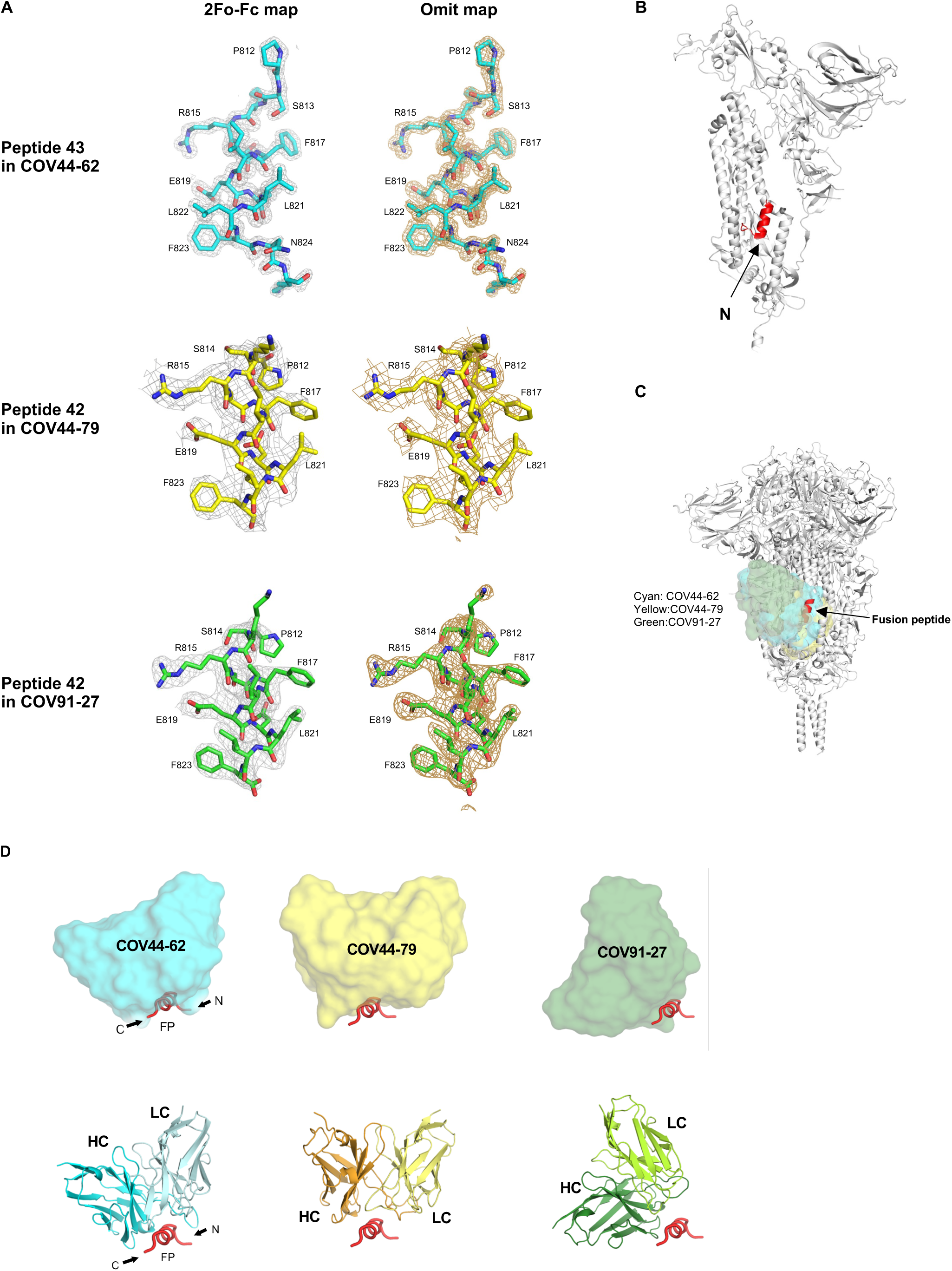
Electron density maps for the fusion peptides and binding angle of COV44-62, COV44- 79, and COV91-27 to the fusion peptide. (**A**) The 2Fo-Fc electron density maps are represented in a gray mesh and contoured at 1.5σ, 0.6σ, and 0.8σ for the fusion peptides (FPs) bound to COV44-62, COV44-79, and COV91-27, respectively. The Fo-Fc unbiased omit electron density maps of the FPs are represented in a brown mesh and contoured at 3.0σ, 1.2σ, and 2.0σ, respectively. (**B**) Location of fusion peptide (in red) on a protomer of the SARS-CoV-2 spike protein (in white). (**C**) Structures of fusion peptides bound with antibodies COV44-62, COV44-79, and COV91-27 were superimposed onto the fusion peptide region (red) of an intact SARS-CoV-2 spike trimer structure in the pre-fusion state (PDB: 6XR8) (*51*). For clarity, only variable domains of the antibodies are shown. The antibodies clash with the spike protein when docked onto their fusion peptide epitopes. Conformational changes or conformational dynamics would be required to fully access the fusion peptide epitope. (**D**) Differences in binding angles of COV44-62, COV44-79, and COV91-27 on interaction with the fusion peptide. The antibodies are aligned on the fusion peptide. The fusion peptides are shown in the same orientation. For clarity, only the variable domains of the antibodies are shown as a molecular surface (top) and in ribbon representation (below).

**Figure S5.**
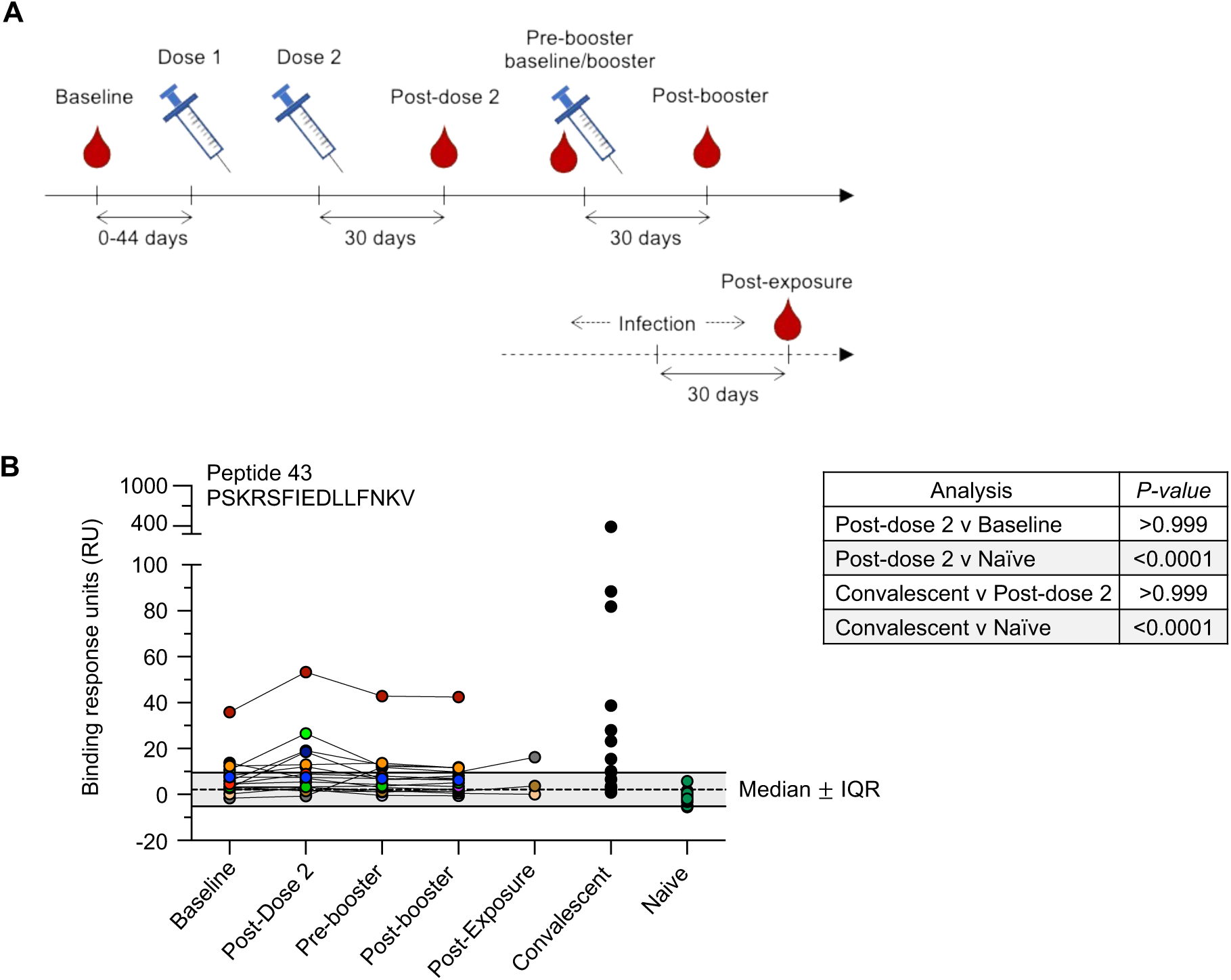
Antibody responses to the fusion peptide after SARS-CoV-2 infection or mRNA-1273 vaccination. (**A**) Vaccination schedule showing plasma or serum collection time points. All individuals were vaccinated and boosted with the Moderna mRNA-1273 vaccine. For three out of 16 individuals, plasma was collected 30 days after documented infection with SARS-CoV-2. (**B**) Circulating IgG reactivity from vaccinated (n=16), convalescent unvaccinated (n=16) and COVID-19 naïve (n=13) individuals to peptide 43, a peptide covering the fusion peptide region just downstream of the S2’ cleavage site. All polyclonal IgG was tested at 100 *μ*g/mL. Background is represented as median ± interquartile range of the donors in the naïve and baseline groups. Statistical analysis was performed using a Kruskal-Wallis test with Dunn’s post-test multiple comparison of the groups shown in the table.

**Figure S6.**
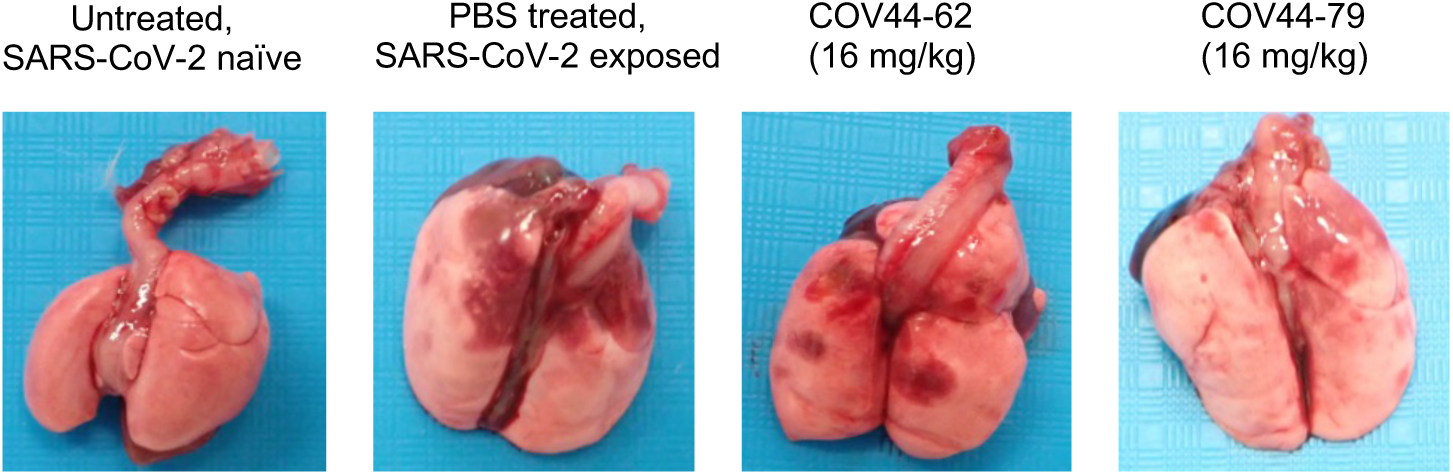
COV44-79 limits lung pathology in SARS-CoV-2 exposed Syrian hamsters. Representative lung pathology of SARS-CoV-2 naïve and SARS-CoV-2 exposed Syrian hamsters from animals that received treatment with COV44-62, COV44-79 or PBS only (control).

**Table S1.**
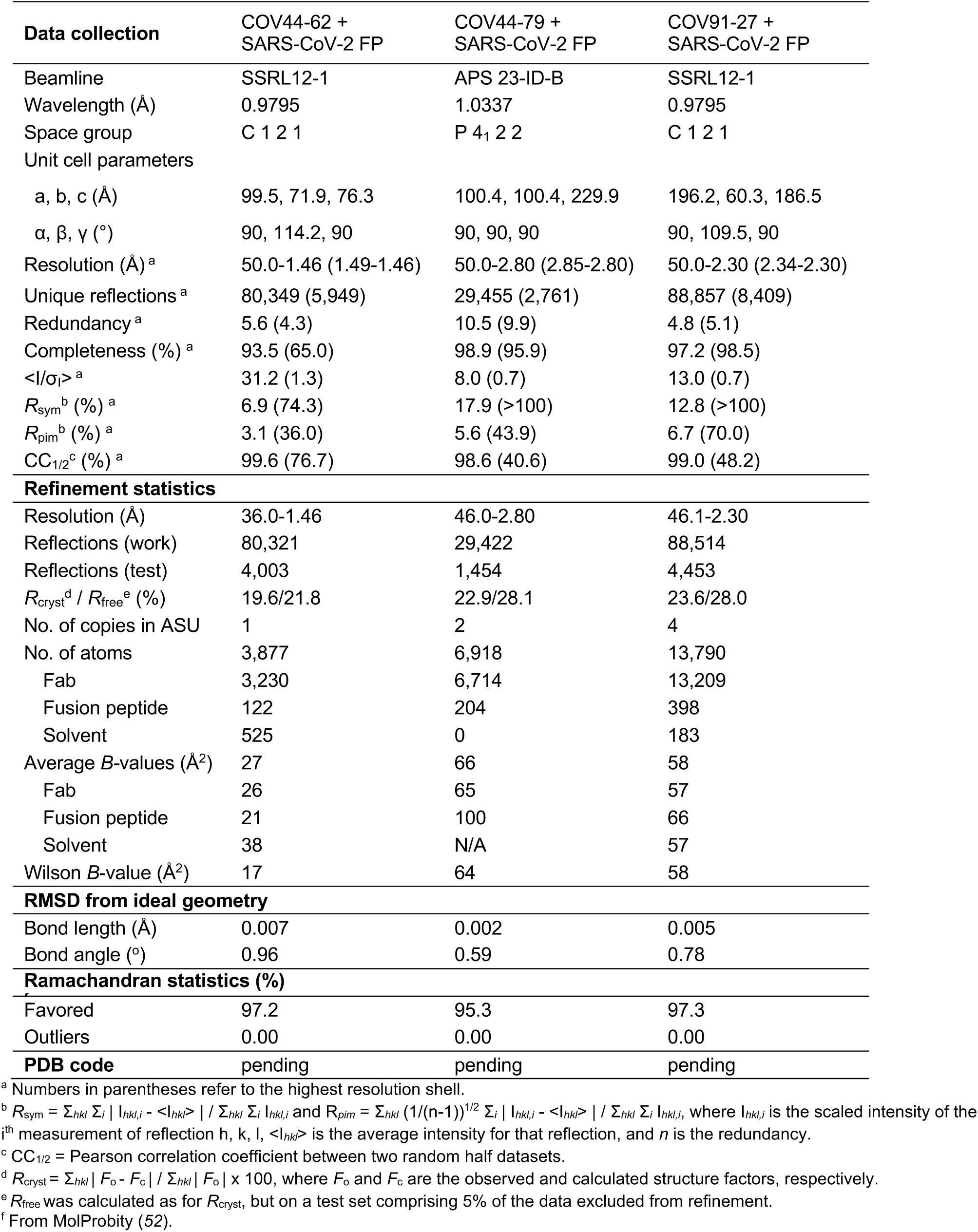
**X-ray data collection and refinement statistics**

## Materials and Methods

### Study participants

For COVID-19 convalescent participants, whole blood and plasma samples were obtained from a previously described cohort (*16*). All participants in this study met the inclusion criteria of ≥ 18 years of age, RT-PCR confirmation of SARS-CoV-2 infection while symptomatic and at least 2 weeks elapsed between their last report of COVID symptoms and the time of blood collection. All participants in the convalescent cohort provided informed consent for their blood products to be used for research purposes by signing the standard New York Blood Center (NYBC) blood donor consent form. Plasma samples from all 142 participants were examined in the initial screen and 19 participants were selected for inclusion in the current study based on plasma IgG reactivity. For participants who received the SARS-CoV-2 mRNA-1273 vaccine (Moderna), whole blood, plasma and serum samples were obtained at the NIH Clinical Research Center in Bethesda, MD under protocols approved by the NIH Institutional Review Board, ClinicalTrials.gov identifiers: NCT00001281 and NCT05078905. All 16 participants in the vaccine study met the inclusion criteria of ≥ 18 years of age, no known history of SARS-CoV-2 infection (based on nucleocapsid antibody responses) and had not previously received a dose of a COVID-19 vaccine at the time of enrollment. Blood samples were serially drawn from all 16 vaccinees: prior to administration of the first mRNA-1273 dose (baseline), 30 days post-administration of the 2^nd^ dose, prior to administration of the 3^rd^ dose (pre-booster baseline) and 30 days after the 3^rd^ dose. For 3 participants, an additional blood draw was collected 30 days post-documented SARS-CoV-2 infection. All participants in the mRNA-1273 cohort provided written informed consent to have their blood products used for research purposes. No randomization or blinding was applied to the analysis of participants’ plasma, serum or PBMC samples, but all samples were anonymized before being used in this study.

### Coronavirus spike proteins

Spike proteins of HCoV-NL63 spike (Sino Biological 40604-V08B,) HCoV-229E spike (Sino Biological 40605-V08B) and HCoV-HKU1 (Sino Biological 40606-V08B) were commercially acquired. The HCoV-OC43 spike, MERS-CoV spike and H1 hemagglutinin proteins were synthesized as previously described (*32*) and kindly gifted by Andrew Ward. Briefly, both ectodomain constructs contain a C-terminal T4 fibritin trimerization domain, an HRV3C cleavage site, an 8× His-Tag, and a Twin-strep-tag for purification. For protein expression, FreeStyle 293- F cells were transected with the spike plasmid of interest, and cultures were harvested at 6-days post-transfection. The spike proteins were purified from the supernatants on complete™ His-Tag Purification Resin before further purification with Superose 6 increase (S6i) 10/300 column (GE Healthcare Biosciences). The H1 HA gene derived from A/Michigan/045/15 H1 was cloned into pcDNA3.4 vector and contains a GCN4 trimerization domain (folded: MKQIEDKIEEILSKIYHIENEIARIKKLIGE) followed by an 8× His-tag and a Twin-Strep-tag at the C-terminus. The HA protein was purified from supernatants using StrepTactin-XT 4FLOW gravity flow columns (IBA Lifesciences). The protein was incubated with BXT elution buffer for 4 h, eluted, and further purified using a Superdex 200 increase 10/300 column (GE Healthcare Biosciences) in TBS buffer. Pre-fusion stabilized constructs for CCoV HuPn-2018 (Accession # QVL91811.1, aa1-1384 with E1140P and E1141P mutations) and PdCoV0081-4 (Accession # MW685622.1, aa1-1092 with E854P and V855P mutations) were synthesized and cloned into pCDNA3.1- vectors (Genscript) with the following C-terminal modifications: T4 fibritin trimerization motif, HRV3C protease cleavage site, poly-GS linker, Avi-tag, and 8× His tag. Plasmids were mixed 1:3 with 1 mg/mL PEIMax, pH 7 (Polysciences) to transfect 293Freestyle cells. At day 3, cells were supplemented with soy hydrolase and glucose. 5-7 days post-transfection, the supernatant was harvested, clarified, and purified using a His-Trap Excel Column (Cytiva).

SARS-CoV-2 NTD, RBD, and spike, as well as SARS-CoV RBD and spike, were expressed and purified as previous described (*16, 33, 34*). Briefly, the SARS-CoV-2 NTD and RBD were cloned into an in-house pFastBac vector. The vector was fused with a gp67 signal peptide and an His6 tag flanking the N- and C-terminus of the NTD and RBD. The recombinant bacmids were obtained from Bac-to-Bac system (Life Technologies). Baculoviruses were produced by transfection of bacmid DNA into Sf9 cells and used to infect High Five cells (Life Technologies) at high (5 to 10) multiplicity of infection (MOI). The supernatant of the infected High Five cells was harvested around 72 hours post-infection at 28°C with shaking at 110 rpm.

For SARS-CoV-2 spike, the SARS-CoV-2 S HexaPro plasmid was a generous gift from Jason McLellan (Addgene plasmid # 154754) (*19*). The spike S2 domain (699 to 1207 with F817P, A892P, A899P, A942P, K986P, V987P) was constructed into phCMV3 vector which contained an N-terminal secreting signal peptide, and C-terminal thrombin cleavage site and His6 tag. The proteins were purified by Ni Sepharose excel resin (Cytiva), and followed by size exclusion chromatography (SEC) in 20 mM Tris buffer with150 mM NaCl, pH7.4.

### Generation of multiplexed CoV antigen beads

Streptavidin beads pre-labelled with individual intensities of PE-channel fluorophore (Spherotech, SVFA-2558-6K and SVFB-2558-6K) or FITC-channel fluorophore (Spherotech, SVFA-2552-6K and SVFA-2552-6K) were incubated with 10 μg/mL each of a subset or all of the following biotinylated antigens: recombinant SARS-CoV-2 spike (Wuhan 1), SARS-CoV-2 RBD, SARS-CoV-2 NTD, SARS-CoV-1 spike and SARS-CoV-1 RBD, MERS-CoV spike, OC43-CoV spike, CCoV-HuPn-2018 spike, pPDCoV-0081-4 spike, HCoV-NL63 spike, HCoV-229E spike, HCoV- HKU1 spike, H1 HA and recombinant CD4 (gifted by Gavin Wright, (*35*)). Antigens acquired in the His-tagged, unbiotinylated form were incubated with 2μg/mL anti-His biotin (Invitrogen, MA1-21315-BTIN) for 20 min at room temperature before being used to label the streptavidin beads. After incubation with the antigens, beads were washed with 0.05% BSA w/v in PBS and incubated with 10 μg/mL of CD4 to block excess streptavidin sites. The blocked beads were washed twice with 0.05% BSA w/v in PBS and mixed to generate multiplexed configurations as required.

### Plasma IgG reactivity to human coronaviruses and donor selection

Multiplexed beads for SARS-CoV-2, SARS-CoV-1, MERS-CoV, HCoV-OC43, HCoV-HKU1, HCoV-229E and HCoV-NL63 spike proteins, as well as CD4 as a negative control, were incubated with donor plasma diluted at 1/50, 1/250 or 1/1250 for 30 min at room temperature, then washed and stained with 2.5 μg/mL goat anti-human IgG Alexa Fluor 647 (Jackson ImmunoResearch, 109-606-170). Samples were acquired on the iQue Screener Plus (Intellicyt) high-throughput flow cytometer and FACS data were analysed with FlowJo (Version 10.8.1., Ashland, OR). Plasma reactivity was analyzed by calculating area under the curve (AUC) for the IgG binding titration curves and reported after correction using the AUC of the negative control CD4 population. All AUC analyses were performed with GraphPad Prism (Version 9.3.1, San Diego, CA) 19 donors were selected for further analysis if positive for plasma reactivity to the spike proteins of SARS-CoV-2 and at least one other beta-coronavirus.

### Memory B cell isolation from PBMCs

Cryopreserved PBMCs were thawed and stained with the following panel: DAPI (BD564907), CD14-BV510 (BioLegend, 301842), CD3-BV510 (BioLegend, 317332), CD56-BV510 (BioLegend, 318340), CD19-ECD (Beckman Coulter, IM2708U), CD21-BV711 (BD, 563163), IgA-Alexa Fluor 647 (Jackson ImmunoResearch, 109-606-011), IgD-PE-Cy7 (BD, 561314), and IgM-PerCP-Cy5.5 (BD, 561285), CD27-Alexa Fluor 488 (BioLegend, 393204) and CD38-APC-Cy7 (BioLegend, 303534). The cells were sorted using the BD FACSAria IIIu in a BSL3 facility and gated on live CD19^+^CD14^-^CD3^-^CD56^-^IgM^-^IgD^-^IgA^+^/IgA^-^ (memory B cells).

### Optofluidic-based isolation of individual cross-reactive antibody secreting B cells

Sorted MBCs (CD19^+^ IgA^+^/IgA^-^) were resuspended with irradiated 3T3-CD40L feeder cells (*36, 37*) in IMDM (Gibco, 31980-030) supplemented with 10% heat-inactivated (HI)-FBS (Gibco, 10438-026), 100 ng/mL IL21 (Gibco, PHC0211), 0.5 μg/mL R848 (Invivogen, tlrl-r848) and 1× Mycozap (Lonza, VZA-2021). The cell suspension was seeded at a density of 50-100 MBCs and 3000 irr-3T3s per well of 384-well plates and co-cultured for 10 days to allow for MBC expansion and activation of Ig secretion. At day 9, culture supernatants were screened for IgA/IgG reactivity against beads coated with 10 μg/mL SARS-CoV-2, SARS-CoV-1, MERS-CoV, OC43-CoV, HKU1-CoV, 229E-CoV and NL63-CoV spike using the iQue screener. On day 10, MBCs from wells of interest were pooled, washed in MACS buffer (PBS supplemented with 0.5% w/v BSA and 2mM EDTA) and approximately 23,000 cells were loaded onto an OptoSelect 11k chip. OEP light cages were applied to sort single B cells into nanoliter-volume pens (nanopens) on the chip and antibody secreting cells were screened for cross-reactivity in a two-step assay. First, 7 μm streptavidin beads (Spherotech, SVP-60-5) coated in tandem with 10 μg/mL MERS-CoV spike and 10 μg/mL OC43-CoV spike were re-suspended in a cocktail of 2.5 μg/mL goat anti-human IgG-Alexa Fluor 647 (Jackson ImmunoResearch, 109-606-170) and goat anti-human IgA-Cy3 (Jackson ImmunoResearch, 109-166-011), and immobilized in the channels of the OptoSelect 11k chip. Binding of secreted antibody to the beads was detected in the CY5 or TRED channels by capturing images at 6 min intervals over a 30 min time course. In the second step of the assay, the MERS/OC43 beads were replaced with 7 μm streptavidin beads coated with 10 μg/mL SARS-CoV-2 spike, and antibody binding was detected as before. OEP light cages were applied to export individual cross-reactive mAb secreting B cells directly into Dynabeads mRNA DIRECT lysis buffer (Life Technologies, 61011) in 96-well plates. Plates were sealed with Microseal foil film (BioRad, MSF1001) and immediately frozen on dry ice before transferring to -80 °C for long-term storage.

### mAb sequence analysis and expression

Heavy and light chain sequences were amplified from single B cell lysates using RT-PCR (*16, 38, 39*) and resolved by Sanger Sequencing (Eurofins and Genewiz). Analyses of the VH and Vλ/Vκ genes, CDR3 sequences, and percentage of somatic mutations were carried out using Geneious Prime (Version 2021.0.3, https://www.geneious.com) and the International Immunogenetics Information System database (IMGT, http://www.imgt.org/) (*40*). Matched pairs of antibody VH and Vλ/Vκ sequences were commercially cloned into plasmids containing an IgG1 or relevant light chain backbone and expressed as recombinant antibody (Genscript). mAbs were also expressed in-house by transient transfection of Expi293 cells (Gibco, A14527) using the ExpiFectamine 293 Transfection Kit (Gibco, A14524) according to manufacturer’s instructions. Recombinant IgG mAbs were purified using HiTrap Protein A columns (Cytiva/GE Healthcare Life Sciences, 17040303).

### HLA typing for donor identification

Several mAbs were isolated from screens that involved pooling of B cells from two different individuals. To identify the source donor of these mAbs, a commercially available ScisGo^®^-HLA- v6 kit (Scisco Genetics Inc., Seattle WA) employing an amplicon-based sequencing by synthesis approach was used to determine HLA types of amplified cDNA from single cell isolates. The approach uses a two-stage amplicon-based PCR for locus amplification and sample barcoding. Although designed for amplification from genomic DNA, a subset of the kit amplicons was functional in amplifying product from cDNA. Briefly, samples were sequentially applied to stage 1 (S1) and stage 2 (S2) PCR amplification according the manufacturer supplied protocol. After amplification, the reactions were combined, purified, and applied to a MiSeq using Illumina Version 2 chemistry with 500-cycle, paired-end sequencing (Illumina, San Diego, CA). Data assembly and analysis were performed using Sciscloud^®^ (Scisco Genetics Inc., Seattle WA) computational tools adapted specifically to assemble HLA genomic sequences derivative from the ScisGo^®^-HLA-v6 kit. Amplified portions of the HLA class I and class II genes were compared with prior typing data allowing the unambiguous identification of corresponding samples. Access to all software for data transfer and analysis was included as a component of the kit and made available through a web browser.

### Recombinant mAb binding to coronavirus antigens

Recombinant mAbs were diluted 4-fold in 0.05% BSA w/v in PBS to generate a 47.7 ng/mL – 200 μg/mL dilution series. Multiplexed antigen-labelled beads were incubated with mAb titrations for 30 min at room temperature, then washed and stained with 2.5 μg/mL goat anti-human IgG Alexa Fluor 647 (Jackson ImmunoResearch, 109-606-170). Samples were acquired on the iQue Screener Plus and FACS data were analysed with FlowJo. Data points from the titration curves were interconnected without logistic regression and AUC analyses were performed with GraphPad Prism and reported after correction using the AUC of the negative control CD4 population.

### Phylogenetic tree generation

Full-length amino acid sequences of SARS-CoV-2 (accession #NC_045512.2), SARS-CoV (accession # AY278741.1), MERS-CoV (accession # NC_019843), HCoV-NL63 (accession #NC_005831.2), HCoV-229E (accession #NC_002645.1), CCoV HuPn-2018 (accession #MW591993.2) and PDCov-0081-4 (accession #MW685622) were aligned using the L-INS-i method of MAFFT version 7. A Neighbor-Joining tree based on the sequence alignments was constructed in MEGA11 with 500 bootstrap resamplings. The phylogenetic tree was visualized in the idol online server (*41*).

### Sequence conservation

Full-length amino acid sequences of spike from SARS-CoV-2 Wuhan-Hu-1 (accession # YP_009724390), SARS-CoV-2 B.1.1.7 (accession # QWE88920), SARS-CoV-2 B.1.351 (accession # QRN78347), SARS-CoV-2 P.1 (accession # QVE55289.1), SARS-CoV-2 B.1.617.2 (accession # QWK65230), SARS-CoV-2 BA.1 (accession # UFO69279.1), SARS-CoV-2 BA.2 (accession # UJE45220.1), Avian Infectious Bronchitis Virus (accession # F4MIW6), BatCoV- HKU3 (accession # Q3LZX1), BatCoV-HKU4 (accession # A3EX94), BatCoV-HKU9 (accession # ABN10911), BatCoV-RaTG13 (accession # QHR63300.2), BatCoV-Rs3667 (accession #AGZ48818), BatCoV-Rs4231 (accession # ATO98157), BatCoV-WIV1 (accession # AGZ48831), WhaleCoV-SW1 (accession # B2BW33), BuCoV-HKU11 (accession # ACJ12044), CCoV-HuPn-2018 (accession # QVL91811), Civet-SARS-CoV-007/2004 (accession # AAU04646), HCoV-229E (accession # NP_073551), HCoV-HKU1 N5 (accession # Q0ZME7), HCoV-NL63 (accession # YP_003767), HCoV-OC43 (accession # P36334), MERS-EMC/2012 (accession #YP_009047204), MuCoV-HKU13-3514 (accession # B6VDY7), Pangolin-CoV-GX/P2V (accession # QIQ54048), PDCoV/Haiti/Human/0081-4/2014 (accession # QWE80492.1), PDCoV/Haiti/Human/0329-4/2015 (accession # QWE80508), SARS-CoV-Urbani (accession # AAP13441.1), Mouse Hepatitis Virus (accession # P11224), ThCoV-HKU12 (accession # B6VDX8), TCoV (accession # B3FHU5), WiCoV-HKU20 (accession # H9BR25) were aligned using MAFFT v7 server using a BLOSUM62 scoring matrix and L-INS-I algorithm. The sequence alignment was used to generate a sequence logo plot using the Weblogo 3.0 server and to color conserved amino acid residues on a pre-fusion stabilized spike protein (PDB 6VSB).

### SARS-CoV-2 spike and S2 subunit epitope binning by SPR

Epitope binning experiments were performed on the Carterra LSA with cross-reactive mAbs coupled to an HC30M chip (Carterra). The chip was conditioned by successive injections of 50 mM NaOH, 500 mM NaCl and 10 mM glycine pH 2, then primed with MES supplemented with 0.05% Tween. To prepare the mAb array, the chip was activated with a 1:1 mixture of 400 mM EDC and 100 mM NHS (ThermoFisher Scientific) followed by direct coupling of 0.1 mg./mL, 1.0 μg/mL and 10 μg/mL of the mAbs in pH 4.5 acetate buffer onto discrete spots on the chip. Excess binding sites on the chip were blocked with 1M ethanolamine, pH 8.5. For pre-mixed binning experiments with trimeric SARS-CoV-2 spike, 20 nM SARS-CoV-2 spike was pre-mixed in a 1:1 volume ratio with 2 μM of each sandwiching antibody and the mAb-spike complexes were then injected onto the array. For classical binning experiments with SARS-CoV-2 S2 domain, 200 nM of monomeric SARS-CoV-2 S2 domain was directly injected onto the chip followed by 10 μg/mL of each sandwiching mAb. The chip was regenerated with 10 mM glycine pH 2.0 after each sandwiching antibody injection. Binning data were analyzed using the Epitope Software (Carterra).

### SARS-CoV-2 S2 binding kinetics with Fab fragments of cross-reactive antibodies

Fab fragments were generated using the Pierce Fab Preparation kit (Thermo Scientific) according to the manufacturer’s instructions. Briefly, 250 mg to 500 mg of mAb diluted in PBS were buffer exchanged into digestion buffer (20 mM cysteine-HCl) using a 7K MWCO Zeba desalting column. Immobilized papain was transferred to a spin-column and equilibrated with 0.5 mL digestion buffer. MAbs were cleaved by papain for 3 hours at 37C in an end-over-end mixer. Digested Fabs were purified using the Protein G HP SpinTrap column (Cytiva) and desalted to remove excess cysteine present following the digestion. Following chip activation, Fabs were coupled to an HC- 30M chip (Carterra) at a concentration of 1.67 μg/mL. Serial dilutions of SARS-CoV-2 Wuhan Hu-1 S2P or S2 were injected onto the chip and kinetics was measured using a 10 min association and 20 min dissociation followed by regeneration with 10 mM glycine pH 2.0. Data were analyzed using the Kinetics Software (Carterra), including bulk shift adjustments to account for changes in buffer refractive index.

### SARS-CoV-2 S2 peptide mapping

Peptides spanning the entire SARS-CoV-2 S2 domain (Ser686- Lys1211, Accession #YP_009724390.1) were commercially synthesized (JPT Peptide Technologies). Each peptide was 15 amino acids in length, overlapped its flanking peptides by 12 residues and carried an N-terminal biotin tag. A further 8 irrelevant oligomers representing random H1 hemagglutinin peptides were included in the panel as negative controls. The lyophilized biotinylated peptides were reconstituted to 1 mg/mL in DMSO, then further diluted in HBSTE supplemented with 0.05% BSA and directly coupled to the streptavidin surface of a SAD200M chip (Carterra) at 0.1 μg/mL and 10 μg/mL. Broadly reactive mAbs were successively injected onto the peptide array at 10 μg/mL and binding data were acquired over a 5 min association phase and 1 min dissociation phase using the Carterra LSA. 3 successive injections of 10 mM glycine pH 2.0 were used to regenerate antibody binding sites on the arrayed peptides after each antibody injection. Data were analyzed using the Epitope Software (Carterra).

### Cell-cell fusion inhibition assay

HeLa cell lines stably expressing either CoV spike proteins or their cognate receptor were generated as previously reported (*15*) and maintained in DMEM + 10% FBS + 1% Pen/Strep + 1% Glutamax. To generate spike expressing cells, HeLa cells were transduced with lentivirus encoding both the nuclear localization signal (NLS)-RFP and a relevant CoV spike protein, stained with corresponding mAbs and sorted to collect the RFP^high^/Spike^high^ population. To generate green fluorescent protein (GFP)-tagged receptor cell lines, HeLa-ACE2 cells were transduced with lentivirus encoding GFP and sorted to collect the GFP^high^/ACE2^high^ population. For fusion inhibition assays, 5000 RFP^+^/Spike^+^ HeLa cells were seeded per well in 96-well plates one day before experiment and cultured overnight. mAbs were added to the wells at a final concentration of 200 μg/mL and cultures were further incubated at 37 °C for 1h. 8,000 GFP^+^/ACE2^+^ HeLa cells were then added to each well and the co-cultures were maintained overnight to allow for syncytia development. After visual confirmation of syncytia under the microscope, the culture medium was replaced with 4% PFA and cells were fixed for 15 min and washed twice with PBS. Fixed cultures were counter-stained with 1 μg/mL Hoechst for 10 min and washed twice with PBS. Images were acquired in A488, A568 and DAPI channels using a BZ-X fluorescence microscope (KEYENCE) and processed using Fiji ImageJ (*42*).

### Shotgun mutagenesis epitope mapping of antibodies by alanine scanning

Epitope mapping was performed essentially as previously described (*43*), using a SARS-CoV-2 (Wuhan Hu-1 strain) S2 subunit shotgun mutagenesis mutation library, made using a full-length expression construct for the SARS-CoV-2 spike glycoprotein. 513 S2 residues (between residues 689 -1247) were mutated individually to alanine, and alanine residues to serine. Mutations were confirmed by DNA sequencing, and clones arrayed in a 384-well plate, one mutant per well. Binding of mAbs to each mutant clone in the alanine scanning library was determined, in duplicate, by high-throughput flow cytometry. A plasmid encoding cDNA for each spike protein mutant was transfected into HEK-293T cells and allowed to express for 22 h. Cells were fixed in 4% (v/v) paraformaldehyde (Electron Microscopy Sciences), and permeabilized with 0.1% (w/v) saponin (Sigma-Aldrich) in PBS before incubation with mAbs diluted in PBS, 10% normal goat serum (Sigma), and 0.1% saponin. mAb screening concentrations were determined using an independent immunofluorescence titration curve against cells expressing wild-type spike protein to ensure that signals were within the linear range of detection. Antibodies were detected using 3.75 μg/mL of Alexa-Fluor-488-conjugated secondary antibodies (Jackson ImmunoResearch) in 10% normal goat serum with 0.1% saponin. Cells were washed three times with PBS/0.1% saponin followed by two washes in PBS, and mean cellular fluorescence was detected using a high-throughput Intellicyt iQue flow cytometer (Sartorius). Antibody reactivity against each mutant spike protein clone was calculated relative to wild-type spike protein reactivity by subtracting the signal from mock-transfected controls and normalizing to the signal from wild-type spike-transfected controls. Mutations within clones were identified as critical to the mAb epitope if they did not support reactivity of the test mAb but supported reactivity of other SARS-CoV-2 antibodies. This counter-screen strategy facilitates the exclusion of spike protein mutants that are locally misfolded or have an expression defect.

### Expression and purification of Fabs for structural studies

Sequences of the variable domains of the heavy chain (VH) and light chain (VL) of COV44-62, COV44-79, and COV91-27 were codon optimized (Genscript) and fused with an N-terminal secreting signal peptide, and a human Fab expressing vector. The three Fabs were expressed by co-transfection of plasmids of heavy and light chain at the ratio of 2:1 in ExpiCHO expression system (Life Technologies) for two weeks according to the manufacturer’s manual. Supernatants were harvested, purified with CaptureSelect CH1-XL resin (Life Technologies), and followed by size exclusion chromatography (SEC) in 20 mM Tris buffer with 150 mM NaCl at pH 7.4 (TBS). Fabs were concentrated to at least 10 mg/ml before crystallization trials.

### Crystallization and structural determination

Fusion peptides were synthesized by GenScript. The complex of each Fab with peptide was formed by mixing each Fab with a 10-fold molar ratio of peptide and incubating overnight at 4°C without further purification. Each complex was adjusted to ∼10 mg/ml in TBS buffer, pH 7.4. The complexes were screened for crystallization on our robotic high-throughput CrystalMation system (Rigaku) at The Scripps Research Institute using the JCSG Core Suite (QIAGEN) as precipitant. Crystallization trials were setup by the vapor diffusion method in sitting drops containing 0.1 μl of protein and 0.1 μl of reservoir solution. The optimized crystallization condition for COV44-62 with fusion peptide was 0.1 M sodium citrate, pH 4, 1 M lithium chloride, and 10% PEG6000. The optimized condition for COV44-79 with fusion peptide was 0.1 M Tris, pH 8.5, 0.01 M nickel (II) chloride, and 20% PEG monomethyl ether 2000 and, for the COV91-27-peptide complex, was 70% 2-methyl-2,4-pentanediol and 0.1M HEPES, pH 7.5. Crystals were harvested on or before day 14 and then soaked in reservoir solution containing 20% (v/v) ethylene glycol as cryoprotectant for COV44-62 and COV44-79 complexes, and 15% (v/v) ethylene glycol for the COV91-27-peptide complex. The harvested crystals were then flash-cooled and stored in liquid nitrogen until data collection. Diffraction data were collected at cryogenic temperature (100 K) at the Stanford Synchrotron Radiation Lightsource on Scripps/Stanford beamline 12-1 with a beam wavelength of 0.97946 Å for the COV44-62 and COV91-27 complexes, and at beamline 23-ID-B of the Argonne Photon Source (APS) with a beam wavelength of 1.033167 Å for the COV44-79 complex. The diffraction data were processed with HKL2000 (*44*). Structures were solved by molecular replacement using Phaser (*45*) with the models generated by Repertoire Builder (https://sysimm.org/rep_builder/) for COV44-62, COV44-79, and COV91-27. Iterative model building and refinement were carried out in Coot (*46*) and PHENIX (*47*), respectively. Buried and accessible surface areas were calculated with PISA (*31*).

### Authentic OC43-CoV-GFP virus propagation and neutralization assay

Rhabdomyosarcoma cells (RD, ATCC CCL-136) were maintained at 37°C and 5% CO_2_ in No-glucose DMEM (Gibco, 11966-025), supplemented with 10% HI-FBS, 4500 mg/mL glucose, 1 mM sodium pyruvate (Gibco, 11360-070), 1 mM HEPES (Gibco, 15630-080) and 50 μg/mL gentamycin (Quality Biological, 120-098-661). RD cells were seeded into a T225cm^2^ flask and cultured to achieve 90% confluency. Cell cultures were inoculated with infectious GFP-tagged HCoV-OC43 at 0.01 MOI in FBS-free, high-glucose DMEM supplemented with 1X GlutaMax (Gibco, 11965-092) and sodium pyruvate. Cultures were maintained at 35°C for 1 h with gentle rocking every 10 min. The inoculum was replaced with prewarmed high glucose DMEM supplemented with 1× Glutamax, 1× non-essential amino acids (Gibco, 12491-015), 2% HI-FBS, 15 mM HEPES and 50 μg/mL gentamicin, and the culture was further maintained for 3-4 days at 35°C and 5-9% CO_2_. To harvest progeny virions, the virus-containing culture media was cleared at 234 × g for 30 min at 4°C, and the cleared supernatant was aliquoted and stored at -80°C. The volume of OC43-GFP virus needed to achieve 75% infection (TCID_75_) of RD cell cultures was determined by endpoint dilution. For neutralization assays, 5 × 10^4^ RD cells were inoculated at TCID_75%_ OC43-GFP virus and incubated for 1h at 35°C. 4-fold serial dilutions (73 ng/mL - 300 μg/mL) of each mAb were incubated with TCID_75_ OC43-GFP virus for 1h at 35°C. 60 μL of mAb-virus mixture was used to inoculate each well containing 5 × 10^4^ RD cells and cultures were incubated for 24 h at 35°C. GFP expression was measured on the iQue Screener Plus and analysed using FlowJo. %Neutralization was determined by (100 × (1-(GFPx Min_GFP_) / (Max_GFP_Min_GFP_)), where uninfected, untreated cells = Min_GFP_ and untreated, infected cells = Max_GFP_.

### Pseudovirus production and neutralization assays (Assay_NIH_)

Codon-optimized cDNA encoding full-length spike from SARS-CoV-2 (GenBank ID: QHD43416.1), SARS-CoV (Urbani_ GenBank: AAP13441.1), MERS-CoV EMC_ GenBank: AFS88936), HCoV-NL63 (GenBank: Q6Q1S2.1) and HCoV-229E (GenBank: AOG74783.1) were synthesized (Genscript), cloned into the mammalian expression vector VRC8400 (*48*) and confirmed by sequencing. These full-length spike plasmids were used for pseudovirus production. Spike-containing lentiviral pseudovirions were produced by co-transfection of packaging plasmid pCMVdR8.2, transducing plasmid pHR’ CMV-Luc, a TMPRSS2 plasmid and full-length spike plasmids from SARS-CoV-2, SARS-CoV, MERS-CoV, HCoV-NL63 and HCoV-229E into 293T cells using Lipofectamine 3000 transfection reagent (ThermoFisher Scientific, Asheville, NC, L3000-001) (*49*). 293 flpin-TMPRSS2-ACE2 cells (provided by Dr. Adrian Creanga, VRC/NIH) were used for SARS-CoV-2, SARS-CoV and hCoV-NL63 while HuH7.5 cells were used for MERS-CoV and hCoV-229E neutralization assay. Cells were plated into 96-well white/black Isoplates (PerkinElmer, Waltham, MA) at 10,000 cells per well the day before infection of pseudovirus. Serial dilutions of mAbs were mixed with titrated pseudovirus, incubated for 45 min at 37°C and added to cells in triplicate. Following 2 h of incubation, wells were replenished with 150 ml of fresh media. Cells were lysed 72 h later, and luciferase activity was measured with Microbeta (Perkin Elmer). 50% neutralization titers (NT50) were calculated using the dose-response-inhibition model with 5-parameter Hill slope equation in GraphPad Prism 9.

### Pseudovirus production and neutralization assays (Assay_Scripps_)

Lentiviral based pseudo-viruses were produced similarly to a previous report (*50*). HEK293T cells were seeded in 6 well plates and grown to ∼80% confluency in DMEM (Lonza, 12-614F) with P/S, glutamine and 10% heat-inactivated FBS. 2.5μg 2^nd^ generation lentivirus backbone plasmid pCMV-dR8.2 dvpr (Addgene, 8455), 2μg pBOBI-FLuc (Addgene, 170674) and 1μg truncated coronavirus spike expressing plasmids (SARS: Addgene #170447; SARS2 #170442; MERS #170448; NL63 #172666; alpha strain #170451; beta #170449; gamma #170450; delta #172320; omicron 180375) were co-transfected in HEK293T with Lipofectamine 2000 (ThermoFisher Scientific, 11668019) to produce single-round infection-competent pseudoviruses. The medium was changed 12-16 hours post transfection. Pseudovirus-containing supernatants were collected 48 and 72 hours post transfection, centrifuged at 1,500 × g for 10 min and the viral titers were measured by luciferase activity in relative light units (RLU) (Bright-Glo Luciferase Assay System, Promega, E2620). The supernatants are aliquoted and stored at -80°C until further use. Pseudotyped viral neutralization assays were performed similar to a previous report (*50*). 20μL pseudovirus supernatant were added into 20 μL serial dilutions of purified antibodies (starting from 100 μg/mL and dilute by 3-fold) in 384-well plates (Corning 3570). The mixture was incubated for one hour at 37°C, after which 5,000 HeLa-hACE2 cells/ well (in 20 μL medium containing 30 μg/ml Dextran) were directly added to the mixture. After incubation at 37°C for 42- 48 h, the medium was aspirated and luciferase activity was measured by adding 25 μL 1× luciferase substrate. Neutralizing activity was calculated by reduction in luciferase activity compared to the virus controls. 50% neutralization titers (NT50) were calculated using the dose-response-inhibition model with 5-parameter Hill slope equation in GraphPad Prism 9.

### Hamsterization of human monoclonal antibodies

Genomes corresponding to the mouse IgG2a heavy and light chains were aligned to the genome assembly MesAur1.0 (GCA_000349665.1) for a female Syrian golden hamster downloaded from Genbank. Hamster genes with the highest homology to the mouse IgG2a heavy chain, lambda and kappa light chains genes were cloned into a pCDNA3.4 vector (Genscript) and expressed in Expi293 cells as described above.

### Syrian hamster efficacy studies

Approximately 5-6 weeks old Golden Syrian hamsters with equal number of males and females were acquired from Envigo (Indianapolis, IN USA). Hamsters were delivered to the IRF facility ten days prior to study onset for acclimation. Individually housed hamsters were assigned to eight groups (n = 12 each) by a statistician according to weight and gender. Study blinding occurred, and animals were randomized to treatment. Antibody products were evaluated in this study with single treatment at a dose of 16 mg/kg (16 mg/kg each of cocktail). PBS-treated (mock-treated) and SARS-CoV-2 naïve (mock-exposed) hamsters were included as controls. Animals were treated with antibody or PBS by intraperitoneal (IP) inoculation 24 hours prior to exposure to 5 log_10_ pfu SARS-CoV-2 (WA01) via intranasal (IN) installation in the prophylaxis study. Animals were weighed prior to study initiation (Day -2) to determine the average weight of each group for calculation of the treatment dose. Following virus inoculation, animals were weighed and observed daily to monitor the clinical signs of disease. Half of the animals in each group were euthanized on day 3 post-infection and the other half of each group was euthanized on day 7. At euthanasia, blood, nasal turbinate and lung tissues were collected for further analysis and necropsies were performed. Lung weight was measured, and the gross pathology scores were assigned by pathologist at the necropsy. None of the animals used in the study reached endpoint criteria that would have required an unscheduled euthanasia.

### Animal ethics statement

Animal research was conducted under an IACUC approved protocols at the Integrated Research Facility, Frederick, Maryland, in compliance with the Animal Welfare Act and other federal statutes and regulations relating to animals and experiments involving animals. The facilities at the IRF where this research was conducted are fully accredited by the Association for Assessment and Accreditation of Laboratory Animal Care, International and adheres to principles stated in the Guide for the Care and Use of Laboratory Animals, National Research Council, 2011.

### Vaccinee and convalescent plasma binding to peptides

Polyclonal IgG antibodies from plasma or sera of vaccinated, convalescent, or naïve donors were purified using the Pierce Protein G Spin Plate (Thermo Scientific). Briefly, plasma or sera were diluted 1:4 in PBS and incubated with Protein G for 30 min, 600 rpm at room temperature. The flowthrough was collected and incubated with the Protein G resin for 15 min to ensure maximal binding. The Protein G resin was washed four times with PBS and the IgG was eluted with Protein G Elution Buffer (Thermo Scientific) and neutralized with 1 M Tris pH 8.0. Purified IgG was desalted using a 40 kDa MWCO Zeba Plate and diluted to 100 μg/mL to assess epitope reactivity.

### Statistical analyses

Neutralization curves were fitted using the dose-response-inhibition model of non-linear regression analysis with 5-parameter Hill slope equation, and 50% antibody neutralization titers (NT50) values were interpolated from the resulting curves. Statistical significance for average body weight was analyzed across the 7-day time-course using a mixed-effects repeated measures model with Dunnett’s post-test multiple comparison. Statistical analyses for hamster clinical and pathology scores were analyzed by Kruskal-Wallis tests with Dunn’s post-test multiple comparison. For all analyses *P < 0.05, **P < 0.01, ***P < 0.001, ****P < 0.0001 and ns, not significant. Descriptive statistics (mean ± SEM or mean ± SD) and statistical analyses were performed using Prism version 9.3.1 (GraphPad). Phylogenetic bootstrap resampling was run with 500 iterations in MEGA11.

## References

1. P. V’Kovski, A. Kratzel, S. Steiner, H. Stalder, V. Thiel, Coronavirus biology and replication: implications for SARS-CoV-2. Nat Rev Microbiol 19, 155–170 (2021).

2. E. Dong, H. Du, L. Gardner, An interactive web-based dashboard to track COVID-19 in real time. Lancet Infect Dis 20, 533–534 (2020).

3. S. Iketani et al., Antibody evasion properties of SARS-CoV-2 Omicron sublineages. Nature, (2022).

4. N. Andrews et al., Covid-19 vaccine effectiveness against the Omicron (B.1.1.529) variant. N Engl J Med, (2022).

5. E. Takashita et al., Efficacy of antibodies and antiviral drugs against Covid-19 Omicron variant. N Engl J Med 386, 995–998 (2022).

6. L. A. VanBlargan et al., An infectious SARS-CoV-2 B.1.1.529 Omicron virus escapes neutralization by therapeutic monoclonal antibodies. Nat Med 28, 490–495 (2022).

7. J. A. Lednicky et al., Independent infections of porcine deltacoronavirus among Haitian children. Nature 600, 133–137 (2021).

8. A. N. Vlasova et al., Novel canine coronavirus isolated from a hospitalized patient with pneumonia in East Malaysia. Clin Infect Dis 74, 446–454 (2022).

9. C. B. Jackson, M. Farzan, B. Chen, H. Choe, Mechanisms of SARS-CoV-2 entry into cells. Nat Rev Mol Cell Biol 23, 3–20 (2022).

10. J. S. Tregoning, K. E. Flight, S. L. Higham, Z. Wang, B. F. Pierce, Progress of the COVID-19 vaccine effort: viruses, vaccines and variants versus efficacy, effectiveness and escape. Nat Rev Immunol 21, 626–636 (2021).

11. T. N. Starr et al., Deep mutational scanning of SARS-CoV-2 receptor binding domain reveals constraints on folding and ACE2 binding. Cell 182, 1295–1310 e1220 (2020).

12. D. Pinto et al., Broad betacoronavirus neutralization by a stem helix-specific human antibody. Science 373, 1109–1116 (2021).

13. M. M. Sauer et al., Structural basis for broad coronavirus neutralization. Nat Struct Mol Biol 28, 478–486 (2021).

14. C. Wang et al., A conserved immunogenic and vulnerable site on the coronavirus spike protein delineated by cross-reactive monoclonal antibodies. Nat Commun 12, 1715 (2021).

15. P. Zhou et al., A human antibody reveals a conserved site on beta-coronavirus spike proteins and confers protection against SARS-CoV-2 infection. *Sci Transl Med*, eabi9215 (2022).

16. H. Cho et al., Bispecific antibodies targeting distinct regions of the spike protein potently neutralize SARS-CoV-2 variants of concern. Sci Transl Med 13, eabj5413 (2021).

17. A. C. Walls et al., Cryo-electron microscopy structure of a coronavirus spike glycoprotein trimer. Nature 531, 114–117 (2016).

18. Y. Yuan et al., Cryo-EM structures of MERS-CoV and SARS-CoV spike glycoproteins reveal the dynamic receptor binding domains. Nat Commun 8, 15092 (2017).

19. C. L. Hsieh et al., Structure-based design of prefusion-stabilized SARS-CoV-2 spikes. Science 369, 1501–1505 (2020).

20. J. F. Chan et al., Simulation of the clinical and pathological manifestations of coronavirus disease 2019 (COVID-19) in a golden Syrian hamster model: implications for disease pathogenesis and transmissibility. Clin Infect Dis 71, 2428–2446 (2020).

21. M. Imai et al., Syrian hamsters as a small animal model for SARS-CoV-2 infection and countermeasure development. Proc Natl Acad Sci U S A 117, 16587–16595 (2020).

22. S. F. Sia et al., Pathogenesis and transmission of SARS-CoV-2 in golden hamsters. Nature 583, 834–838 (2020).

23. R. Kong et al., Fusion peptide of HIV-1 as a site of vulnerability to neutralizing antibody. Science 352, 828–833 (2016).

24. K. Xu et al., Epitope-based vaccine design yields fusion peptide-directed antibodies that neutralize diverse strains of HIV-1. Nat Med 24, 857–867 (2018).

25. R. Kong et al., Antibody lineages with vaccine-induced antigen-binding hotspots develop broad HIV neutralization. Cell 178, 567–584 e519 (2019).

26. M. A. Tortorici et al., Broad sarbecovirus neutralization by a human monoclonal antibody. Nature 597, 103–108 (2021).

27. C. H. Shen et al., VRC34-antibody lineage development reveals how a required rare mutation shapes the maturation of a broad HIV-neutralizing lineage. Cell Host Microbe 27, 531–543 e536 (2020).

28. S. Kratochvil et al., Vaccination in a humanized mouse model elicits highly protective PfCSP-targeting anti-malarial antibodies. Immunity 54, 2859–2876 e2857 (2021).

29. A. Wellner et al., Rapid generation of potent antibodies by autonomous hypermutation in yeast. Nat Chem Biol 17, 1057–1064 (2021).

30. C. M. Poh et al., Two linear epitopes on the SARS-CoV-2 spike protein that elicit neutralising antibodies in COVID-19 patients. Nat Commun 11, 2806 (2020).

31. E. Krissinel, K. Henrick, Inference of macromolecular assemblies from crystalline state. J Mol Biol 372, 774–797 (2007).

32. S. Bangaru et al., Structural mapping of antibody landscapes to human betacoronavirus spike proteins. bioRxiv, 2021.2009.2030.462459 (2021).

33. H. Lv et al., Homologous and heterologous serological response to the N-terminal domain of SARS-CoV-2 in humans and mice. Eur J Immunol 51, 2296–2305 (2021).

34. M. Yuan et al., A highly conserved cryptic epitope in the receptor binding domains of SARS-CoV-2 and SARS-CoV. Science 368, 630–633 (2020).

35. C. Crosnier et al., A library of functional recombinant cell-surface and secreted *P. falciparum* merozoite proteins. Mol Cell Proteomics 12, 3976–3986 (2013).

36. J. Huang et al., Isolation of human monoclonal antibodies from peripheral blood B cells. Nat Protoc 8, 1907–1915 (2013).

37. S. Moir et al., CD40-mediated induction of CD4 and CXCR4 on B lymphocytes correlates with restricted susceptibility to human immunodeficiency virus type 1 infection: potential role of B lymphocytes as a viral reservoir. J Virol 73, 7972–7980 (1999).

38. L. T. Wang et al., A potent anti-malarial human monoclonal antibody targets circumsporozoite protein minor repeats and neutralizes sporozoites in the liver. Immunity 53, 733–744 e738 (2020).

39. T. Tiller et al., Efficient generation of monoclonal antibodies from single human B cells by single cell RT-PCR and expression vector cloning. J Immunol Methods 329, 112–124 (2008).

40. M. P. Lefranc, Immunoglobulin and T cell receptor genes: IMGT((R)) and the birth and rise of Immunoinformatics. Front Immunol 5, 22 (2014).

41. I. Letunic, P. Bork, Interactive Tree Of Life (iTOL) v5: an online tool for phylogenetic tree display and annotation. Nucleic Acids Res 49, W293–W296 (2021).

42. J. Schindelin et al., Fiji: an open-source platform for biological-image analysis. Nat Methods 9, 676–682 (2012).

43. E. Davidson, B. J. Doranz, A high-throughput shotgun mutagenesis approach to mapping B-cell antibody epitopes. Immunology 143, 13–20 (2014).

44. Z. Otwinowski, W. Minor, Processing of X-ray diffraction data collected in oscillation mode. Methods Enzymol 276, 307–326 (1997).

45. A. J. McCoy et al., Phaser crystallographic software. J Appl Crystallogr 40, 658–674 (2007).

46. P. Emsley, B. Lohkamp, W. G. Scott, K. Cowtan, Features and development of Coot. Acta Crystallogr D Biol Crystallogr 66, 486–501 (2010).

47. P. D. Adams et al., PHENIX: a comprehensive Python-based system for macromolecular structure solution. Acta Crystallogr D Biol Crystallogr 66, 213–221 (2010).

48. D. H. Barouch et al., A human T-cell leukemia virus type 1 regulatory element enhances the immunogenicity of human immunodeficiency virus type 1 DNA vaccines in mice and nonhuman primates. J Virol 79, 8828–8834 (2005).

49. L. Naldini, U. Blomer, F. H. Gage, D. Trono, I. M. Verma, Efficient transfer, integration, and sustained long-term expression of the transgene in adult rat brains injected with a lentiviral vector. Proc Natl Acad Sci U S A 93, 11382–11388 (1996).

50. T. F. Rogers et al., Isolation of potent SARS-CoV-2 neutralizing antibodies and protection from disease in a small animal model. Science 369, 956–963 (2020).

51. Y. Cai et al., Distinct conformational states of SARS-CoV-2 spike protein. Science 369, 1586–1592 (2020).

52. V. B. Chen et al., MolProbity: all-atom structure validation for macromolecular crystallography. Acta Crystallogr D Biol Crystallogr 66, 12–21 (2010).

